# C-BERST: Defining subnuclear proteomic landscapes at genomic elements with dCas9-APEX2

**DOI:** 10.1101/171819

**Authors:** Xin D. Gao, Li-Chun Tu, Aamir Mir, Tomás Rodriguez, Yuehe Ding, John Leszyk, Job Dekker, Scott A. Shaffer, Lihua Julie Zhu, Scot A. Wolfe, Erik J. Sontheimer

## Abstract

Mapping proteomic composition at distinct genomic loci and subnuclear landmarks in living cells has been a long-standing challenge. Here we report that d<u>C</u>as9-APEX2 <u>B</u>iotinylation at genomic <u>E</u>lements by <u>R</u>estricted <u>S</u>patial <u>T</u>agging (C-BERST) allows the rapid, unbiased mapping of proteomes near defined genomic loci, as demonstrated for telomeres and centromeres. By combining the spatially restricted enzymatic tagging enabled by APEX2 with programmable DNA targeting by dCas9, C-BERST has successfully identified nearly 50% of known telomere-associated factors and many known centromere-associated factors. We also identified and validated SLX4IP and RPA3 as telomeric factors, confirming C-BERST’s utility as a discovery platform. C-BERST enables the rapid, high-throughput identification of proteins associated with specific sequences, facilitating annotation of these factors and their roles in nuclear and chromosome biology.

---

Three-dimensional organization of chromosomes is being defined at ever-increasing resolution through the use of Hi-C and related high-throughput methods^1^. Genome organization can also be analyzed in live cells by fluorescence imaging, especially via fluorescent protein (FP) fusions to nuclease-dead *Streptococcus pyogenes* Cas9 (dSpyCas9), which can be directed to nearly any genomic region via single-guide RNAs (sgRNAs)^2^. It has proven more difficult to map subnuclear proteomes onto 3-D genome landscapes in a comprehensive manner that avoids demanding fractionation protocols, specific DNA-associated protein fusions [e.g. in proximity-dependent biotin identification (BioID^3^)], or validated antibodies. dSpyCas9 has enabled a BioID-derived subnuclear proteomic technique called CasID^4^, in which biotin ligase (BirA*) fusion to dSpyCas9 allows proteins associated with specific genomic regions to be biotinylated on neighboring, exposed lysine residues in live cells. Streptavidin affinity selection and liquid chromatography/tandem mass spectrometry (LC-MS/MS) is then used to identify the tagged proteins. However, this approach is relatively inefficient and usually involves long (18-24h) labeling times, limiting the time resolution of dynamic processes. Similar considerations apply to the recently reported CAPTURE approach for subnuclear proteomics^5^, in which dSpyCas9 itself is biotinylated by BirA* and used as an affinity handle.

Engineered ascorbate peroxidase (APEX2) has been used for an alternative live-cell biotinylation strategy called spatially restricted enzymatic tagging (SRET)^6,7^. In this approach, APEX2 is fused to a localized protein of interest, and cells are then treated with biotin-phenol (BP) and H_2_O_2_, generating a localized (within a ~20nm radius) burst of diffusible but rapidly quenched biotin-phenoxyl radicals. These products react with electron-rich amino acid side chains (e.g. Tyr, Trp, His and Cys), leading to covalent biotinylation of proteins in the vicinity of the localized APEX2, thus allowing subsequent identification by streptavidin selection and LC-MS/MS. Notably, this subcellular tagging method is extremely efficient (1 min H_2_O_2_ treatment), allowing temporal control over the labeling process. Based in part on the success of dSpyCas9-FP fusions in enabling subnuclear imaging in living cells, we reasoned that a dSpyCas9 derivative that emits radicals rather than photons could be used for subnuclear proteomic analyses in a manner that overcomes the unfavorable kinetics and high background of CasID. Here we use dSpyCas9-APEX2 fusions in the development of C-BERST (**Fig. 1a**) for genomic element-specific profiling of subnuclear proteomes in live cells.

**Figure 1.**
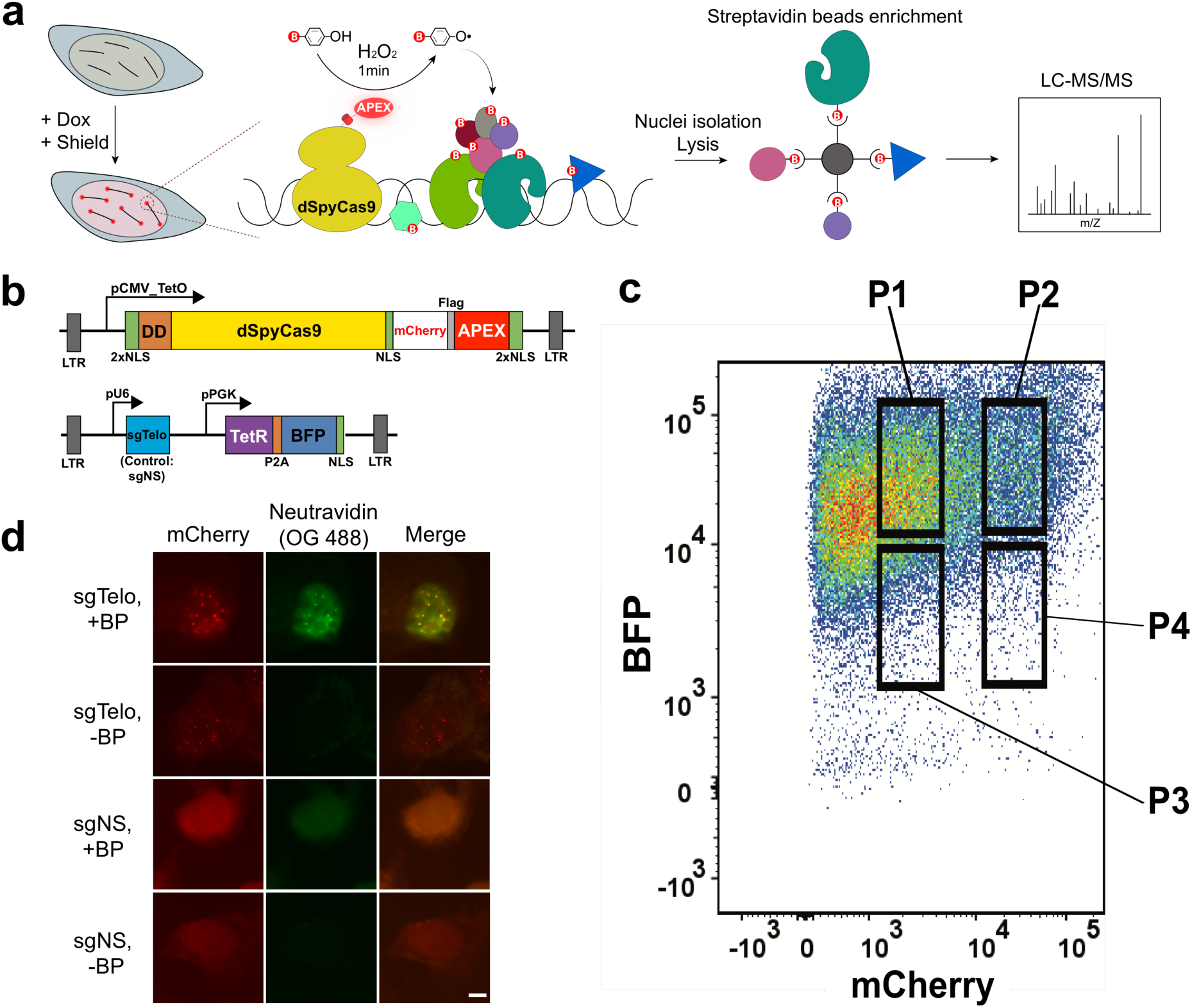
Using C-BERST to biotinylate telomere-associated proteins in living human cells. **(a)** Diagram of the C-BERST workflow. U2OS cells stably expressing sgRNA and inducible dSpyCas9-APEX2 are generated by lentiviral transduction. Following dox and Shield1 induction (21 h), cells are incubated with biotin-phenol (BP, 30 min) and then H_2_O_2_ (1 min) to activate a burst of biotin-phenoxyl radical generation by dSpyCas9-APEX2, leading to proximity-labeling of nearby proteins. Following quenching, nuclei isolation and protein extraction, biotinylated proteins are enriched by streptavidin selection and analyzed by LC-MS/MS. **(b)** The dSpyCas9-mCherry-APEX2 and sgRNA lentiviral expression constructs. Top: dSpyCas9-mCherry-APEX2 under the control of the pCMV_TetO inducible promoter. The mCherry fusion is included to enable quantification of dSpyCas9 expression level as well as its subcellular localization. NLS, nuclear localization signal; LTR, long terminal repeat; DD, Shield1-repressible degradation domain. Bottom: sgRNA/TetR/BFP expression construct. pU6, U6 promoter; pPGK, PGK promoter; sgTelo, telomere-targeting sgRNA; sgNS, non-specific sgRNA; tetR, tet repressor; P2A, 2A self-cleaving peptide; BFP, blue fluorescent protein. **(c)** FACS sorting of untransduced (blue) and mCherry-and BFP-positive cells (red). The P1 population corresponds to high BFP (as a surrogate for sgRNA and TetR) and low mCherry expression, providing optimal signal-to-noise ratio to maximize the fraction of telomere-localized dSpyCas9-mCherry-APEX2. **(d)** Fluorescence imaging of dSpyCas9-mCherry-APEX2 labeling in cells. Stable sgTelo and sgNS cells were labeled live as described in **(a)** or were only supplemented with H_2_O_2_ as a no-labeling control. Cells were then fixed and stained with neutravidin conjugated with OG488 to visualize biotinylated proteins. dCas9-mCherry-APEX2 localization are indicated by mCherry fluorescence. Scale bar, 5 pm.

## Results

### C-BERST design and workflow

To develop and validate this method, we sought genomic elements that are associated with a well-defined suite of known protein factors, and that can be bound with dSpyCas9 with high efficiency and specificity using an established sgRNA. For this purpose, we chose to target telomeres in human U2OS cells. As with ~10-15% of cancer cell types, U2OS cells rely on alternative lengthening of telomeres (ALT) pathways to maintain telomere length without telomerase activation^8^. Cohorts of proteins associated with telomeres in ALT+ cells are well-characterized, and they map to key pathways such as homologous recombination (HR) and break-induced telomere synthesis^9^. Furthermore, an sgRNA (sgTelo) has already been established for efficient telomere association of dSpyCas9^10^’ ^11^.

We transduced U2OS cells with a lentiviral vector expressing dSpyCas9 under the control of a tet-on CMV promoter and fused to five nuclear localization signals (NLSs), a ligand-tunable degradation domain (DD)^12^, mCherry, and APEX2 (**Fig. 1b**). Maximal expression of this dSpyCas9-mCherry-APEX2 fusion protein requires not only doxycycline (dox) but also the

Shield1 ligand to inactivate the DD in a dose-dependent fashion^13^. This combination allows precise control over dSpyCas9-mCherry-APEX2 protein levels for optimal signal-to-noise levels. mCherry-positive cells [collected by fluorescence-activated cell sorting (FACS)] were then transduced with a separate lentiviral vector that included an sgRNA construct (driven by the U6 promoter) as well as a blue fluorescent protein (BFP) construct that also expresses the TetR repressor (**Fig. 1b**). In one version of this construct the sgRNA cassette encodes sgTelo (for labeling telomeres), and in the other it encodes a non-specific sgRNA (sgNS) that is complementary to a bacteriophage-derived sequence that is absent from the human genome^14^. After 21h of dox and Shield1 induction, we again used fluorescence-activated sorting (FACS) to sort four distinct BFP/mCherry double-positive cell populations (P1-P4) that correlate with different expression levels of dSpyCas9-mCherry-APEX2 and BFP (as a surrogate for sgRNA and TetR) (**Fig. 1c** and **Supplementary Fig. 1**). We reasoned that our signal-to-noise ratio of telomeric vs. non-telomeric biotinylation would be maximized when sgTelo levels are saturating, and when dSpyCas9-mCherry-APEX2 levels are limiting (relative to potential genomic binding sites). Both conditions are expected to favor maximal partitioning of the sgRNA-programmed dSpyCas9-mCherry-APEX2 into the desired telomere-associated state, with as little unlocalized or mislocalized fusion protein as possible. Visual inspection by fluorescence microscopy (**Supplementary Fig. 2**) confirmed that the sgTelo P1 cell population (with higher BFP expression and lower mCherry expression, about 20% of the total sorted cells) exhibited the most robust mCherry-labelled telomeric foci (as determined by colocalization with anti-TERF2IP immunofluorescence) with the lowest amount of nucleolar or diffuse nucleoplasmic background^10^. We therefore used this population in our subsequent experiments.

To assess the distribution of biotinylated proteins in the nucleus, we used avidin-conjugated Oregon Green 488 (OG 488) to probe sgTelo and sgNS cells after BP and H_2_O_2_ treatment. We found that biotinylated proteins were strongly enriched at telomeric foci in sgTelo cells, whereas labeling was diffuse in sgNS cells (**Fig. 1d**). Efficient labeling was BP‐ and H_2_O_2_-dependent. We also analyzed genome-wide dCas9-mCherry-APEX2 binding via anti-mCherry chromatin immunoprecipitation and sequencing (ChIP-seq). Nearly 60% of total trimmed reads from sgTelo cells contained at least one (TTAGGG)4 sequence (the minimal length of telomeric repeats with full complementarity to sgTelo). However, such (TTAGGG)4-containing reads comprised <0.5% of total trimmed reads from either sgNS or untransduced U2OS cells (**Supplementary Fig. 3**). These experiments indicate that sgTelo-guided dCas9-mCherry-APEX2 targets telomeres and enables restricted biotinylation of endogenous proteins near these chromosomal elements.

### Label-free profiling of telomere-associated proteomes using C-BERST

For proteomic analysis, we induced APEX2-catalyzed biotinylation with BP and H_2_O_2_ in the sgTelo and sgNS P1-sorted cells (~6 x 10^7^ cells for each guide), and also included an sgTelo control in which the H_2_O_2_ was omitted. APEX2-catalyzed biotinylation of nucleoplasmic proteins in the sgNS control sample serves as a reference, permitting an assessment of the telomere specificity of labeling in the sgTelo sample. Nuclei were isolated from treated cells (to reduce cytoplasmic background), and nuclear proteins were then extracted. Recovered proteins (50 μg) were subjected to western blot analysis using streptavidin-conjugated horseradish peroxidase (streptavidin-HRP) (**Fig. 2a**), as well as total protein visualization by Coomassie staining (**Fig. 2b**). Samples were also probed with anti-mCherry antibodies (to detect dSpyCas9-mCherry-APEX2) and HDAC1 (as a loading control) (**Fig. 2a**, bottom). Biotinylated proteins were readily detected in both sgTelo and sgNS samples, but were largely absent in the ‐H_2_O_2_ control (**Fig. 2a**). In the sgNS sample, anti-mCherry and streptavidin-HRP signals were less intense in comparison with the sgTelo sample, indicating that dSpyCas9-mCherry-APEX2 accumulation and activity are lower in the former. Biotinylated proteins were then isolated using streptavidin beads and analyzed by SDS-PAGE and silver staining (**Fig. 2c**). Aside from the ~75kDa endogenously biotinylated proteins routinely detected in SRET-labeled samples^6,7^, only background levels of proteins were detected in the no-H_2_O_2_ control sample, indicating successful purification. All three samples were subjected to in-gel trypsin digestion followed by LC-MS/MS to identify the biotin-labeled proteins. These analyses were done with two biological replicates prepared on different days.

**Figure 2.**
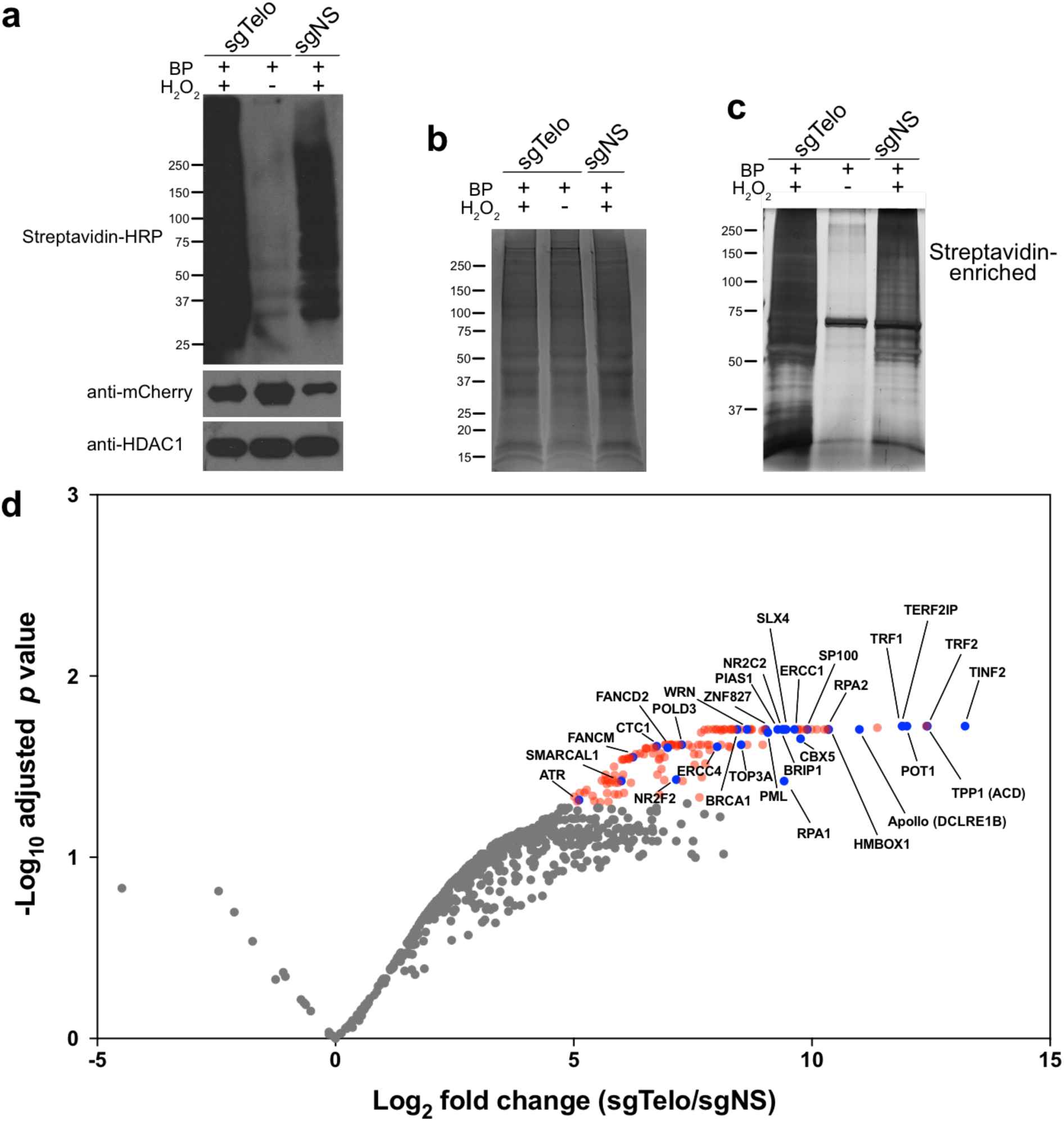
Successful capture of telomere-associated proteins in living human cells by C-BERST using Label Free Quantification (LFQ). **(a)** Top: Western blot analysis of dSpyCas9-mCherry-APEX2 biotinylation, as detected by streptavidin-HRP. sgRNAs, BP treatment, and H_2_O_2_ treatment are indicated at the top of each lane. Anti-mCherry was used to detect dSpyCas9-mCherry-APEX2 (middle), and anti-HDAC1 was used as a loading control (bottom). **(b)** Coomassie-stained SDS-PAGE of total protein from isolated nuclei following biotin labeling. **(c)** Silver-stained SDS-PAGE of biotin-labeled proteins enriched with streptavidin-coated beads. In **a-c**, the mobilities of protein markers (in kDa) are indicated on the left of each panel. **(d)** Volcano plot of C-BERST-labeled, telomere-associated proteins in U2OS cells. Intensity-based absolute quantification (iBAQ) values from the MS analyses were calculated for each identified protein for all three samples (sgTelo + H_2_O_2_, sgTelo - H_2_O_2_, and sgNS + H_2_O_2_). 143′proteins (indicated by blue and red) are statistically enriched [Benjamini-Hochberg (BH)-adjusted *p* value < 0.05] in the sgTelo + H_2_O_2_ sample, relative to both control samples. The 30 proteins indicated by blue dots (with identities provided) are previously defined as either telomere-associated proteins or ALT pathway components. These include all six shelterin components.

The two sgTelo replicates yielded at least three peptides from 930 and 851 proteins, >85% of which (792) were detected in both (**Supplementary Table 1**). For these 792 proteins, we used intensity-based absolute quantification (iBAQ) values [a label-free quantification (LFQ) proteomic approach] to determine the degree of enrichment in the sgTelo sample relative to the sgNS sample. Some of these 792 proteins (104 in the first replicate, and 340 in the second) yielded no spectra whatsoever in the corresponding sgNS sample, consistent with sgTelo specificity. In those cases, to avoid infinitely large sgTelo/sgNS enrichment scores, we assigned those proteins the smallest non-zero iBAQ value from the proteins positively identified in that sgNS dataset. The sgTelo/sgNS iBAQ ratios were then analyzed by moderated t-test, yielding 143 proteins whose enrichment in sgTelo was statistically significant [Benjamini-Hochberg (BH)-adjusted *p* < 0.05] [**Fig. 2d** (red and blue dots) and **Supplementary Table 1**]. Strikingly, the six subunits of the shelterin complex (a telomere-binding complex that protects ends from chromosome fusion^15^) were among the seven most significantly enriched proteins (**Fig. 2d** and **Supplementary Table 1**). Another highly enriched protein was Apollo, a 5’**→**3’ exonuclease that interacts with the shelterin component TRF2 and functions in the ALT pathway^16^. Overall, among the 143 most significantly sgTelo-enriched proteins (**Supplementary Table 1**), 30 have been reported previously to be associated with telomeres or linked to telomere function (**Supplementary Table 2**). These results indicate that validated telomeric proteins can be identified rapidly and efficiently by C-BERST.

### Ratiometric proteomics enhances the sensitivity and specificity of C-BERST

To further improve our assessments of differential C-BERST biotinylation with specific vs. non-specific sgRNAs, we used a more quantitative proteomic approach enabled by stable isotope labeling with amino acids in cell culture (SILAC) to analyze telomere-associated proteomes. sgTelo/dCas9-mCherry-APEX2 cells were cultured in heavy-isotope medium, sgNS/dCas9-mCherry-APEX2 cells were cultured in medium-isotope medium, and untransduced U2OS cells were cultured in light-isotope medium, each for at least 5 passages to allow sufficient incorporation of isotope-labeled arginine and lysine (**Supplementary Fig. 4a**). We then induced dCas9-mCherry-APEX2 expression by dox and Shield1 for 21 hours, with comparable accumulation with either sgTelo or sgNS (**Supplementary Fig. 4b**). Biotinylation and cell lysis were then performed as described above, except that equal amounts (~1 mg, measured by Pierce BCA Protein Assay Kit) of protein lysates from heavy, medium, and light samples (H:M:L = 1:1:1) were mixed before streptavidin affinity purification for three-state SILAC^6^. 913 proteins were identified in both the heavy and medium samples, and 885 of these were also detectable in the light (no-APEX2 background) sample. Using significance (BH-adjusted *p* < 0.01) and enrichment ([log_2_ fold change (FC) ⩾ 2.5]) cut-offs that were even more stringent than those used for the label-free analysis (**Fig. 2d**), we identified 55 proteins that are strongly enriched in the sgTelo sample relative to sgNS (H/M) (**Fig. 3** and **Supplementary Table 3**). Among these 55 proteomic hits, 34 are known telomere-associated factors, including all six shelterin components as well as subunits from 5 other complexes that are known to contribute to ALT-associated pathways or processes (**Supplementary Fig. 5**). All but one of the 55 H/M-enriched proteins (BARD1) were also strongly enriched (log2 FC ⩾ 1) in H/L ratio, indicating that background detection in the absence of dCas9-mCherry-APEX2 biotinylation was minimal. Gene ontology (GO) analysis of the 55 H/M-enriched C-BERST hits reveals strong functional associations with terms such as telomere maintenance, DNA replication, DNA repair, and homologous recombination, all of which are important for ALT pathways^8^ (**Fig. 4a**).

**Figure 3.**
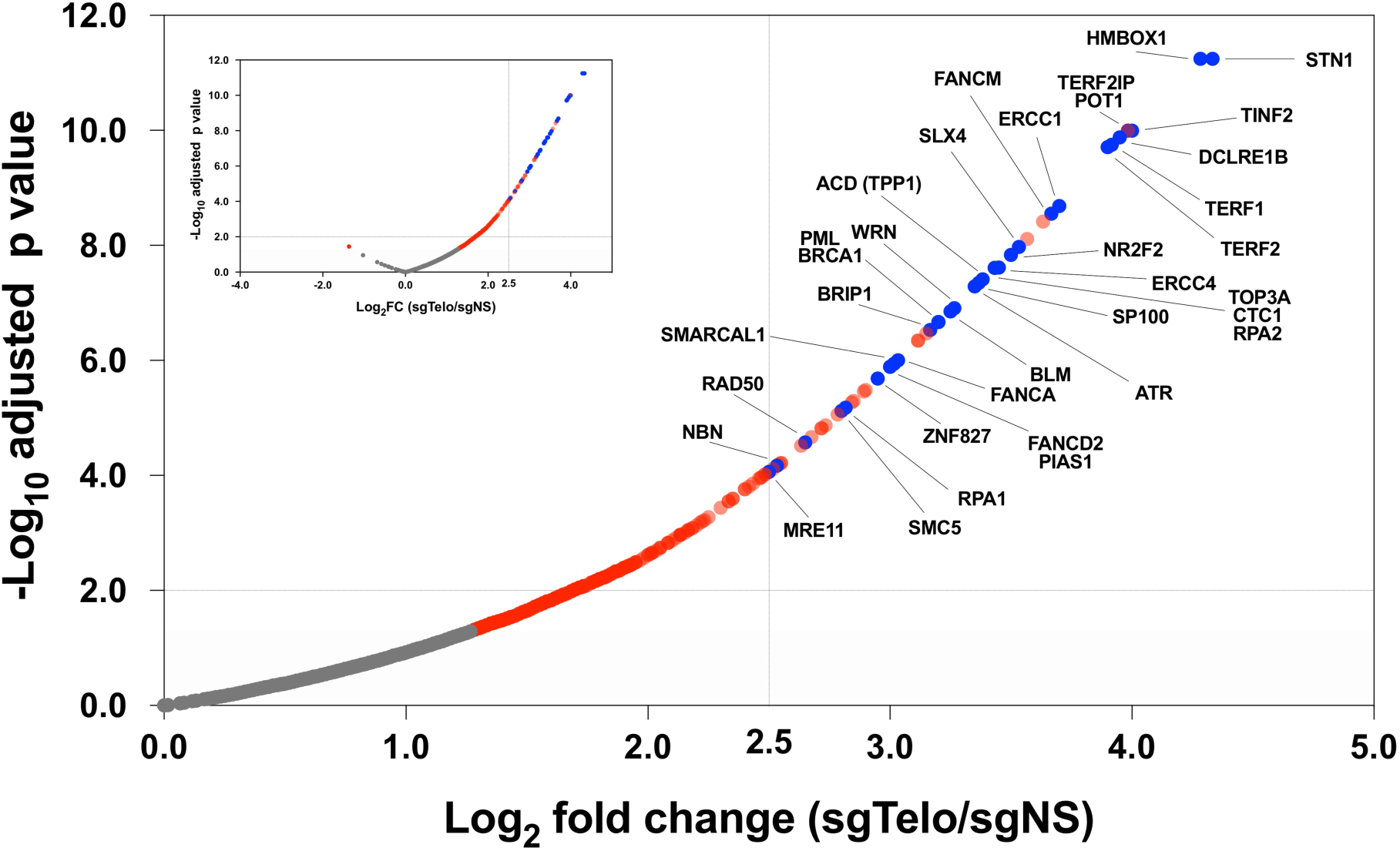
Ratiometric C-BERST tagging strategy improves telomere-associated proteome identification. Volcano plot of C-BERST-labeled, telomere-associated proteins in U2OS cells. For each protein, the H/M SILAC ratio reflects the enrichment of identified proteins in sgTelo vs. sgNS cells. 359 proteins (indicated by blue and red) are statistically enriched [Benjamini-Hochberg (BH)-adjusted *p* value < 0.05] in the sgTelo, relative to sgNS controls (indicated by red dots). 55 proteins fall within the cut-off based on FDR and enrichment level (BH-adjusted *p* value < 0.01 and log_2_ fold change > 2.5). The 34 proteins indicated by blue dots (with identities provided) are previously defined as either telomere-associated proteins or ALT pathway components. The volcano plot shows 97.1% of identified proteins (inset shows all proteins, including the few with SILAC H/M log2 ratios < 0).

Telomere-associated proteomes from ALT+ cell lines have been defined previously by TRF1-BirA* BioID in U2OS cells^17^, and by biochemical purification [proteomics of isolated chromatin segments (PICh)] from WI38-VA13 cells^18^. Protein identifications from these analyses, as well as from our C-BERST SILAC dataset, were examined for overlap as depicted in the Venn diagram shown in **Fig. 4b**. Of the 55 proteins identified by C-BERST, 32 (~58%) were also detected by one or both of the other methods [23 by BioID ( *p* = 7.29 x 10^-32^) 27 by PICh ( *p* = 2.04 x 10^-50^), and 18 by both]. The remaining 23 proteins that were uniquely detected by C-BERST include seven known telomeric/ALT factors (ATR, CTC1, FANCA, FANCD2, FANCM, SMC5, and WRN). Of the 18 proteins detected by all three approaches, 17 are known telomere-related factors. The remaining consensus hit [SLX4-interacting protein (SLX4IP)] was not previously validated as telomeric in the BioID and PICh studies; nonetheless its identification by all three proteomic approaches strongly suggests that it has a previously unappreciated role in telomere function or maintenance (**Fig. 4c**). Such a role could be related to that of its binding partner SLX4 in the resolution of telomere recombination intermediates^19^.

**Figure 4.**
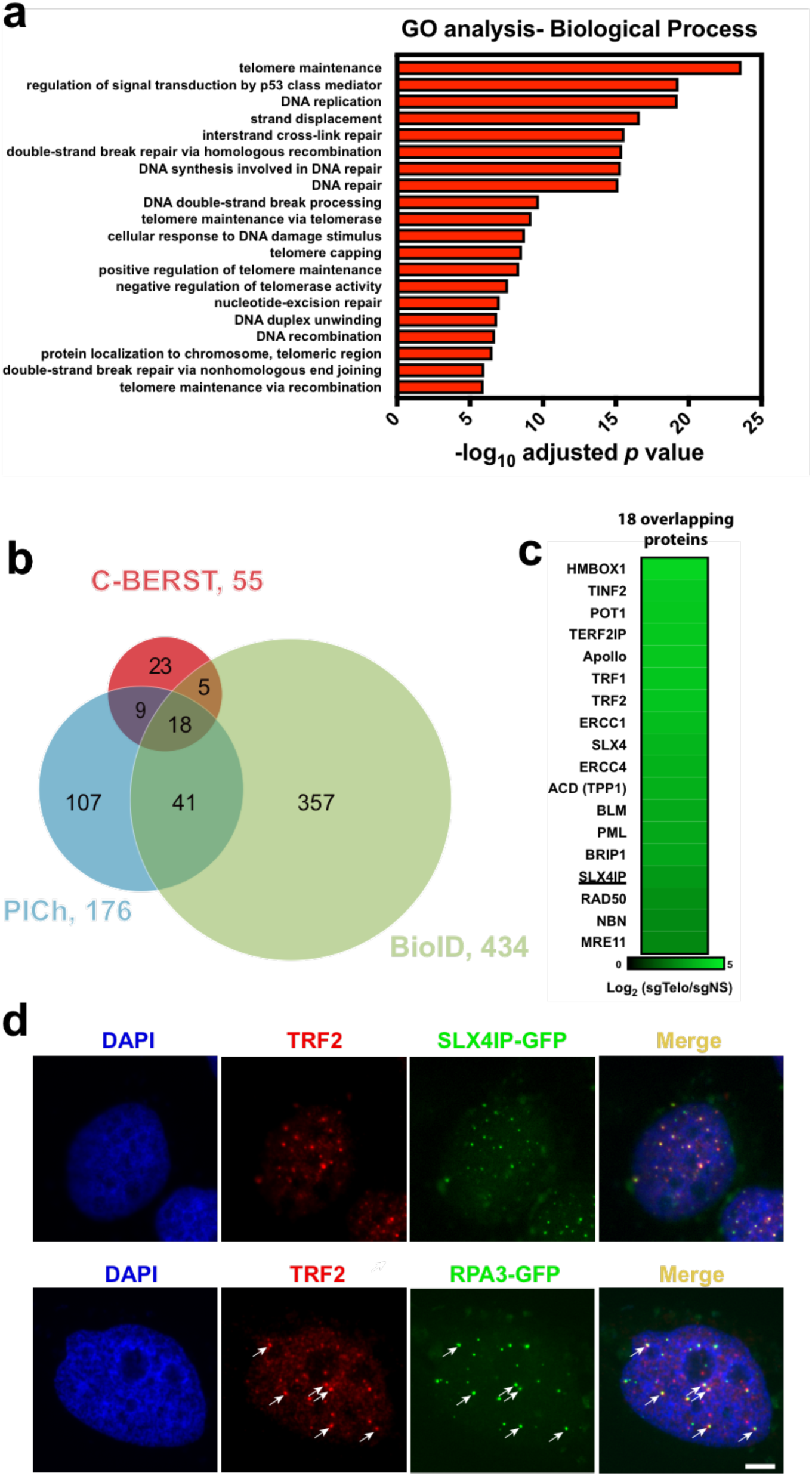
Comparison of the C-BERST telomere-associated proteome (based on the SILAC dataset) with other approaches, and validation of novel telomeric factors. **(a)** Gene Ontology-Biological Process (GO-BP) analysis on the 55 telomeric/ALT proteins identified by C-BERST. The x-axis is the ‐log_10_ *p* value (BH-adjusted) for the C-BERST-detected proteins associated with each GO term given on the left. The 20 most statistically significant GO terms are displayed. **(b)** Venn diagram of statistically enriched (BH-adjusted *p* value < 0.01) telomeric proteins from ALT human cells, as detected by C-BERST (red), PICh (purple), and TRF1-BirA* BioID (green). 32 proteins from the C-BERST proteome were also detected by PICh, BioID, or both. **(c)** A heat map of the C-BERST log2 fold-change enrichment scores for the 18 telomeric proteins identified by all three proteomic approaches from (b). All 18 proteins are highly enriched in the C-BERST telomere proteome. The one underlined protein (SLX4IP) has not been reported previously (to our knowledge) as telomere-or ALT-associated. **(d)** Colocalization of turboGFP-tagged SLX4IP and RPA3′with telomeric marker protein TRF2. 0.3′x 10^5^ U2OS cells were transiently transfected with 100ng SLX4IP-GFP expression plasmid or 50ng RPA3-GFP expression plasmid. Cells were then fixed and incubated with TRF2 primary antibody and secondary antibody conjugated with Alexa Fluor 647. DAPI stained cells were imaged (n ≥ 20 cells examined). Scale bar, 5 pm.

To validate the ability of C-BERST to identify novel or provisional telomeric or ALT-related proteins, we used independent methods to assess telomere colocalization of SLX4IP, as well as a factor (RPA3) that was detected by C-BERST [ranked 44^th^ (log_2_FC) among the 55 enriched proteins] but missed by BioID and PICh. We transiently transfected U2OS cells with turboGFP-tagged SLX4IP, and then analyzed cells for its appearance in endogenous TRF2-colocalizing foci, as indicated by anti-TRF2 immunofluorescence (**Fig 4d**). SLX4IP-GFP signal was evident in some but not all TRF2 foci, confirming that it localizes to a subset of telomeres. This conclusion was further corroborated by immunofluorescence detection of endogenous SLX4IP, which again revealed TRF2 colocalization (**Supplementary Fig. 6**). We also transfected U2OS cells with RPA3-GFP, and again we detected it in a subset of TRF2 foci (**Fig 4d**). Due to high background staining by the commercial anti-RPA3 antibody, we were unable to analyze TRF2 colocalization by endogenous RPA3, but western analyses confirmed that the RPA3-GFP fusion protein (like the anti-SLX4IP fusion protein) was expressed at or below the levels of the endogenous protein under the conditions used for fluorescence microscopy (**Supplementary Fig. 7**). RPA3 is a subunit of the RPA complex, which has known functions in ALT pathways^20^; in addition, other RPA subunits (RPA1 and RPA2) were enriched in PICh^18^ as well as C-BERST. Therefore the partial telomeric localization of RPA3 is not altogether surprising, despite the fact that it was not detected by either PICh or BioID. The incomplete overlap for SLX4IP and RPA3 with the telomeric marker is consistent with previously defined non-telomeric functions for both SLX4IP^21^ and RPA3^22^ and indicate that C-BERST is sufficiently sensitive and specific to detect telomere-associated factors even with proteins that are only partially (or perhaps transiently) telomeric.

### C-BERST subnuclear proteomics at centromeres

To extend our subnuclear proteomic approach to other genomic elements, we targeted dCas9-mCherry-APEX2 to centromeric alpha-satellite arrays in U2OS cells, and then used C-BERST to profile the protein components of alphoid chromatin (**Figure 5a**) using a similar pipeline as that described above for telomeres. The human alpha satellite proteome from K562 cells has been analyzed previously using the PICh-related protocol known as HyCCAPP (hybridization capture of chromatin-associated proteins for proteomics), again providing a basis for comparison. Via live-cell imaging (**Figure 5b**) and western blotting (**Supplementary Fig. 8**), we have confirmed dCas9-mCherry-APEX2 inducible expression, specific centromere targeting^23^, and biotinylation. We used SILAC proteomic analysis (heavy, sgAlpha; medium, sgNS; light, untransduced U2OS cells lacking dCas9-mCherry-APEX2) and identified 1268 proteins (**Supplementary Table 4**) from each of two biological replicates (based upon detection in at least two of three technical replicates of each biological replicate). Among these 1268 proteins, 460 were enriched to a statistically significant extent (log_2_FC ≥ 2.5 and *p* < 0.01) in the sgAlpha vs. sgNS samples (H/M). We have identified four highly enriched subunits of the CENP-A nucleosome-associated complex^24^ (CENP-C, ‐M, ‐N, and ‐T), and one subunit (CENP-P) of the CENP-A distal complex^24^. We also identified nearly all CENP-A loading factors^25^, including HJURP, Mis18α, Mis18β, and MIS18BP1, among the enriched proteins. Many other known centromere-associated proteins were also identified as enriched in sgAlpha including CENP-B, ‐F, ‐I, and ‐L, as well as KNL1. Additionally, we identified three subunits of the chromosome passenger complex [INCENP, CDCA8, and Aurora kinase B (AURKB)], which has been suggested to localize to the centromere during mitosis^26^. We also found Fanconi anemia pathway proteins such as FANCM, which has been implicated in promoting centromere stability^27^. There are 31 overlapping enriched proteins ( *p* = 3.84 x 10^-38^) between C-BERST and HyCCAPP, despite the use of different cancer cell lines in the respective studies (**Supplementary Fig. 9a**). C-BERST uniquely captured multiple known centromeric factors including CENP-F and ATR, which were recently reported to localize to centromeres and engage RPA-coated R loops^28^. GO analysis of the 460 C-BERST centromeric hits reveals strong functional associations with terms such as DNA repair, DNA replication, sister chromatid cohesion, double-strand break repair by homologous recombination, mitotic nuclear division, and cell division, all of which are related to centromere maintenance or function^8^ (**Supplementary Fig. 9b**).

**Figure 5.**
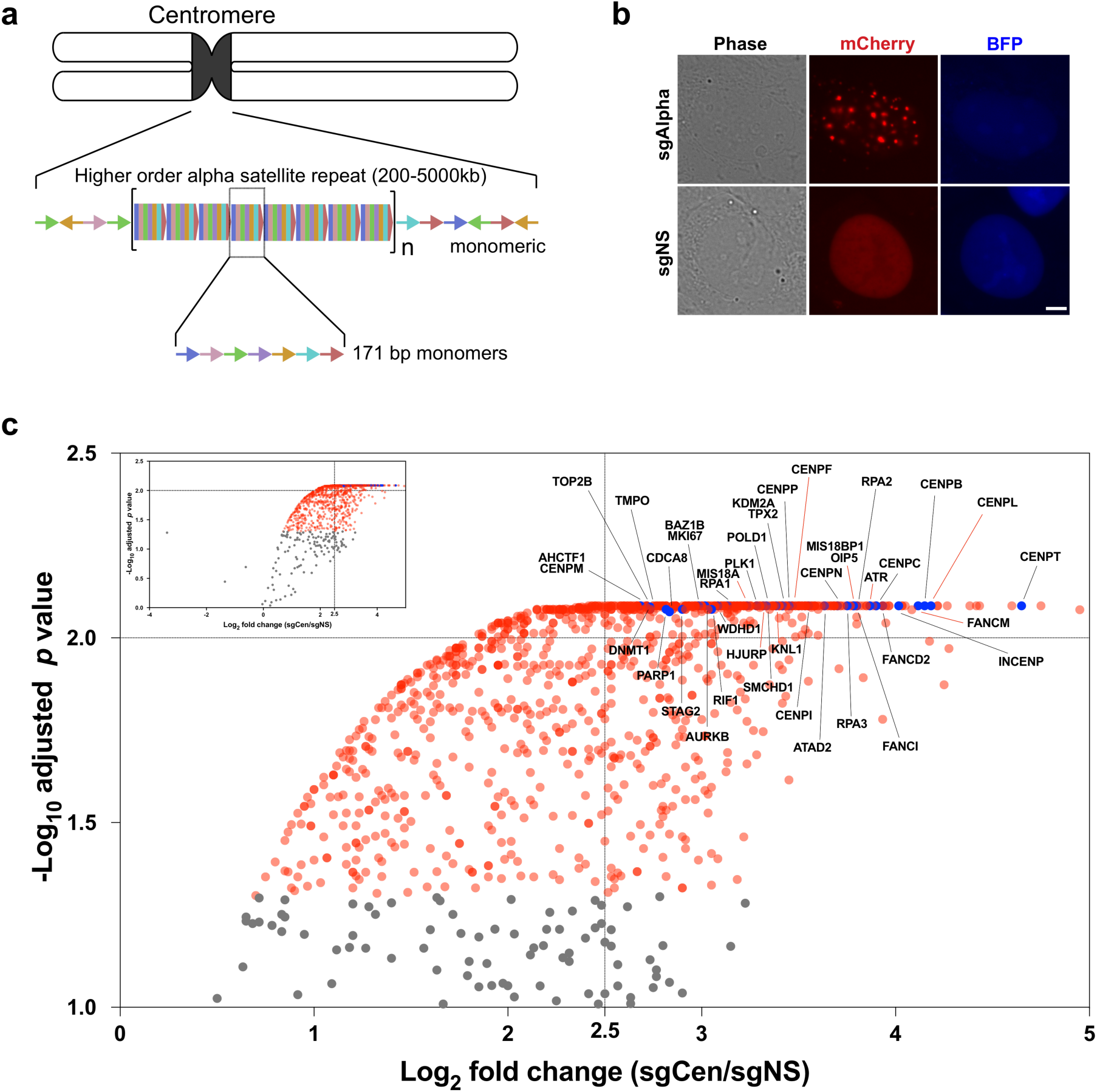
Successful capture of alpha satellite associated proteomes in live human cells by C-BERST. **(a)** Schematic diagram of alpha satellite repeat position and arrangement at a centromere. **(b)** Live-cell imaging of centromere localization by dSpyCas9-mCherry-APEX2 in U2OS cells, using the P1-sorted population defined by the FACS workflow in **Fig. 1c**. dSpyCas9-mCherry-APEX2 exhibited centromeric foci with sgAlpha but not with sgNS. Scale bar, 5 pm. **(c)** Ratiometric C-BERST (using SILAC) was used to profile the alpha satellite associated proteome. A volcano plot of C-BERST-labeled, centromere-associated proteins in U2OS cells is shown. For each protein, the H/M SILAC ratio reflects the enrichment of identified proteins in sgAlpha vs. sgNS cells. 1134 proteins (indicated by blue and red dots) are statistically enriched [Benjamini-Hochberg (BH)-adjusted *p* value < 0.05] in the sgAlpha sample, relative to sgNS controls. 460 proteins fall within the cut-off based on FDR and enrichment level (BH-adjusted *p* value < 0.01 and log_2_ fold change ≥ 2.5). The 40 proteins indicated by blue dots (with identities provided) were previously defined as either centromere-associated proteins (**Supplementary Table 5**) or were reported as components of the HyCCAPP centromere proteome (see text). The nine known centromere-associated proteins indicated by red lines are uniquely captured by C-BERST. The volcano plot shows 96.2% of identified proteins (inset shows all proteins, including the few with SILAC H/M ratio < 0 and ‐log_10_ adjusted *p* value < 1).

Our generation of both telomeric and centromeric C-BERST datasets affords the opportunity to compare SILAC-based protein enrichment at these two chromosomal landmarks. Of the 55 and 460 C-BERST enriched proteins at ALT + telomeres and centromeres, respectively, 36 were identified in both ( *p* = 1.31 x 10^-57^) (**Supplementary Fig. 10**). Significant GO terms for these 36 overlapping proteins include DNA replication, regulation of signal transduction by p53 class mediator, strand displacement, double strand break repair via homologous recombination, and DNA repair, each of which would be expected for both categories of chromosomal elements. Significantly, all CENP factors were found among the 424 non-overlapping proteins from the sgAlpha centromeric dataset. Conversely, the 19 telomere-specific hits (enriched only with sgTelo and not sgAlpha) include five of the six shelterin subunits (TRF1, TERF2IP, TIN2, POT1, TPP1); intriguingly, the sixth (TRF2) has previously been reported to associate with CENP-F^29^. These results provide strong evidence that C-BERST successfully measures subnuclear protein enrichment at distinct chromosomal elements.

## Discussion

We demonstrate that C-BERST successfully maps subnuclear proteomes associated with genomic landmarks. Using the extensively investigated ALT telomeric proteome as an initial benchmark for our ratiometric implementation of C-BERST, we recover approximately 44% of known ALT-associated proteins (34 of 78, **Supplementary Tables 2 and 3**) as strongly enriched at telomeres, in addition to factors involved in all reported biological processes that contribute to ALT. Combining C-BERST with SILAC made it possible to set very high enrichment cut-offs (BH-adjusted *p* < 0.01, log_2_FC ⩾ 2.5]) while still retaining excellent representation of the known telomeric or centromeric factors that we employed as benchmarks; lower thresholds can be set where appropriate to cast a wider net for previously unknown factors. We used fluorescence microscopy to validate a strongly enriched C-BERST hit (SLX4IP) whose telomeric localization had not been previously confirmed, and another (RPA3) that was missed by previous subnuclear proteomic approaches such as BioID^17^ and PICh^18^. Importantly, we extended C-BERST to a second category of genomic elements (alpha satellite centromeric sequences) and identified a distinct set of enriched factors that included many known centromeric factors, and that largely excluded known telomere-specific factors (including those enriched by telomeric C-BERST). These results provide strong indications that C-BERST can successfully profile subnuclear proteomes based upon proximity to specific classes of genomic sequence. C-BERST is also compatible with other ratiometric approaches such as tandem mass tagging (TMT)^30^. Although ratiometric approaches increase the sensitivity and quantitative rigor of C-BERST enrichment, more economical label-free quantitation can also be used successfully with C-BERST, as we showed for telomeres.

By combining the flexibility of RNA-guided dSpyCas9 genome binding with the efficiency and rapid kinetics of APEX2-catalyzed biotinylation, C-BERST promises to extend the unbiased definition of subnuclear proteomes to many other genomic elements, and to a range of dynamic processes (e.g. cellular differentiation, responses to extracellular stimuli, and cell cycle progression) that occur too rapidly to analyze via the longer labeling procedures often necessary for CasID^4^, CAPTURE^5^, and related BirA*-based approaches. Furthermore, we found 40-60 million cells to provide a sufficient sample size for telomeric and centromeric C-BERST, in contrast to the >10^9^ cells reported as inputs for CasID, CAPTURE, and PICh^4,5,18^. The ability to apply C-BERST to smaller populations of cells provides an obvious cost savings, and may also be important when available cell numbers are limiting due to their proliferation properties or challenges in their manipulation. Finally, C-BERST and BirA*-based methods favor biotinylation of distinct sets of proteins by virtue of their different labeling specificities (lysines for BirA*, and predominantly tyrosines for C-BERST); using these approaches in tandem would likely diminish the number of false negatives resulting from inefficient labeling due to differences in the surface-accessible amino acid distribution or the suitability of certain peptides for MS analysis.

Importantly, C-BERST promises to augment and extend Hi-C and related methods by linking conformationally important cis-elements with the factors that associate with them. Guide RNA multiplexing should enable the extension of C-BERST subnuclear proteomics to singlecopy, non-repetitive loci. In the meantime, many types of repetitive elements within the genome, like telomeres and centromeres, play critically important roles in chromosome maintenance and function in ways that depend upon their associated proteins; C-BERST provides an unbiased method for sampling subnuclear, locus-specific proteomics at these elements to define protein factors that are critical to their functions.

## METHODS

Methods, data files, and any associated references are available in the online version of the paper. *Note: Any Supplemental and Source Data files are available in the online version of the paper*.

## ACKNOWLEDGEMENTS

We are grateful to all members of the Sontheimer, Wolfe and Dekker labs for advice and discussions, Tom Fazzio, Samyabrata Bhaduri and Michael Green for helpful feedback, Hanhui Ma, Tong Wu, David Grünwald, and Thoru Pederson for reagents, the Flow Cytometry Core Facility at UMass Medical School for cell sorting, and Lingji Zhu for assistance with figure preparation. This work was supported by 4D Nucleome grant U54 DK107980 from the National Institutes of Health to J.D., S.A.W. and E.J.S.

## AUTHOR CONTRIBUTIONS

X.D.G. and E.J.S. conceived the study. X.D.G., L.-C.T., J.D., S.A.W., and E.J.S. designed experiments. X.D.G. and T.R. performed C-BERST and ChIP-seq experiments, and J.L. conducted mass spectrometry procedures. X.D.G and L.-C.T. processed the fluorescence images, A.M. processed flow cytometry data, X.D.G., Y.D., and J.L. processed mass spectrometry data, and L.J.Z. conducted statistical analyses. All co-authors interpreted the data. X.D.G. and E.J.S wrote the manuscript, and all authors revised and edited the manuscript.

## COMPETING FINANCIAL INTERESTS

The authors declare no competing financial interests.

Reprints and permissions information is available online at http://www.nature.com/reprints/index.html

## ONLINE METHODS

### Construction of C-BERST plasmids

The Shield1‐ and doxycycline-inducible dSpyCas9-mCherry-APEX2 construct was made by subcloning Flag-APEX2 from Flag-APEX2-NES (Addgene 49386) into DD-dSpyCas9-mCherry^13^ using the pHAGE backbone. Two additional NLSs (SV40 and nucleoplasmin NLS) were inserted at each terminus to improve nuclear localization. The sequence of the final plasmid is provided in the **Supplementary Note**. The sgTelo-encoding construct was created by replacing the C3-guide RNA sequence (pCMV_C3-sgRNA_2XBroccoli/pPGK_TetR_P2A_BFP) with sgTelo sequences (using a plasmid provided by Hanhui Ma and Thoru Pederson). Non-specific sgRNA (sgNS)^14^ and sgAlpha were constructed similarly. SLX4IP-turboGFP plasmid was obtained from OriGene (catalog number: RG220896). RPA3-turboGFP was made by replacing the SLX4IP coding sequence with the human RPA3 coding sequence.

### Cell culture and cell line construction

Human U2OS cells obtained from Thoru Pederson’s lab (originally obtained from ATCC) were cultured in Dulbecco-modified Eagle’s Minimum Essential Medium (DMEM; Life Technologies) supplemented with 10% (vol/vol) FBS (Sigma). Lentiviral transduction was as described^13^. Six-fold higher titers of sgRNA-encoding lentiviruses were used for transduction relative to dSpyCas9-APEX2 lentivirus. Stably transduced cells were grown under the same conditions as the parental U2OS cells.

### Flow cytometry

One day before performing FACS, dox (Sigma; 2 μg/ml) and Shield1 (Clontech; 250 nM) were added to the media. ~2 x 10^6^ cells expressing dSpyCas9-mCherry-APEX2 and BFP sgRNA were selected by FACSAria cell sorter or analyzed with MacsQuant^®^ VYB. Both instruments are equipped with 405‐ and 561-nm excitation lasers, and the emission signals were detected by using filters at 450/50 nm (wavelength/bandwidth) for BFP, and 610/20 nm (FACSAria) or 615/20nm (MacsQuant) for mCherry. Bulk population and single cells (**Supplementary Fig. 1b**) were sorted into plates containing 1% GlutaMAX, 20% FBS, and 1% penicillin/streptomycin in DMEM medium.

### Fluorescence microscopy

U2OS cells expressing sgRNA were seeded onto 170 μm, 35 x 10 mm glass-bottom dishes (Eppendorf) supplemented with dox and Shield1 21 hours before imaging. Live cells were imaged with a Leica DMi8 microscope equipped with a Hamamatsu camera (C11440-22CU), a 63x oil objective lens, and Microsystems software (LASX). Further imaging processing was done with MetaMorph (Molecular Devices). Image contrast was set to ease visualization of cell, foci and nucleoplasmic background.

### Immunofluorescence microscopy

Cells for immunofluorescence microscopy were grown on glass coverslips. The transfected cells or normal cells were fixed for 15 minutes in 2% paraformaldehyde in PHEM [0.05 M PIPES/0.05 M HEPES (pH 7.4), 0.01 M EGTA, 0.01 M MgCh_2_], followed by a 2-min. extraction with 0.1% Triton X-100 in PHEM. After PBS washes, the cells were blocked by 1% BSA/1X TBST at 4°C overnight. Cells were first incubated with primary antibodies for two hours at room temperature and washed three times with blocking solution (10 minutes/wash). Cells were then incubated with secondary antibodies for one hour at room temperature, followed by another three blocking solution washes and two PBS washes^31^. Cells were mounted with ProLong antifade and visualized by fluorescence microscopy as described above. Neutravidin conjugated with OG488 experiment was described previously^6^.

### C-BERST biotinylation protocol

Six 15cm plates of U2OS cells (~6 x 10^7^) expressing specific (sgTelo or sgAlpha) or nonspecific (sgNS) sgRNAs were used in this assay. Dox (2 μg/ml) and Shield1 (250 nM) were added 21 hours before biotinylation. Cells were then incubated with 500 μM biotin-phenol (BP) (Adipogen) for 30 minutes at 37°C. 1 mM H_2_O_2_ was then added to initiate of biotinylation for 1 minute on a horizontal shaker at room temperature. Six 15cm plates of sgTelo-or sgAlpha-expressing cells were treated in parallel, but without H_2_O_2_ addition, as a negative control. Quencher solution (5 mM trolox, 10 mM sodium ascorbate, and 10 mM sodium azide) was added to stop the reaction, and cells were washed five times (three quencher washes and two DPBS washes) to continue the quench and to remove excess BP.

### Enrichment of biotinylated proteins

Cells were scraped off the plates and used for the preparation of isolated nuclei^32^. Nuclei were washed with DPBS before lysis. RIPA lysis buffer [50 mM Tris-HCl (pH 7.5), 150 mM NaCl, 0.125% SDS, 0.125% sodium deoxycholate and 1% Triton X-100 in Millipore water) with 1x freshly supplemented Halt Protease Inhibitor were used to lyse the cells for 10 minutes on ice. Cell lysates in 1.5 ml Eppendorf tubes were sonicated for 15 minutes with a Diagenode Bioruptor with 30s on/off cycles at high intensity. Cell lysates were clarified by centrifugation at 13,000 rpm for 10 minutes. Clarified protein samples (~3.5 mg) were subjected to 400 μl Dynabeads MyOne Streptavidin T1 affinity purification overnight at 4°C. Each bead sample was washed with a series of buffers to remove non-specifically bound proteins: twice with RIPA lysis buffer, once with 1 M KCl, once with 0.1 M Na_2_CO_3_, once with 2 M urea in 10mM Tris-HCl, pH 8.0, and twice with RIPA lysis buffer. Proteins were eluted in 70 μl 3x protein loading buffer supplemented with 2 mM biotin and 20 mM DTT with heating for 10min at 95°C^6^. 50 μl eluents were loaded and run on a 4-12% SDS-PAGE gel (Bio-Rad) and run approximately 1cm off the loading well for in-gel digestion and LC-MS/MS analysis. The gel-fractionated sample used for LC-MS/MS (see below) corresponded to proteins from ~4 x 10^7^ cells.

### Western blotting

Protein concentrations of the cell lysates were determined by BCA assay (Thermo). 50 ug of each sample was mixed with protein loading buffer, boiled, and separated in SDS-PAGE gels. Proteins were transferred to PVDF membrane (Millipore), and blotted with Streptavidin-HRP (Thermo), or with anti-mCherry (Abcam) or anti-HDAC1 (Bethyl) antibodies. Additional details of the anti-SLX4IP and anti-RPA3 western analyses are described in the figure legends.

### mCherry affinity purification of dSpyCas9-mCherry-APEX2 captured DNA and sequencing

1 × 10^7^ U2OS cells stably expressing dCas9-mCherry-APEX2 transduced with sequence-targeting or non-specific sgRNAs were washed with PBS, fixed with 1% formaldehyde for 10 min and quenched with 0.125 M glycine for 5 min. Cells were harvested using a plate scraper and lysed in RIPA cell lysis buffer [50 mM Tris-HCl (pH 7.5), 150 mM NaCl, 0.125% SDS, 0.125% sodium deoxycholate and 1% Triton X-100 in Millipore water] with 1x freshly supplemented Halt Protease Inhibitor for 10 minutes on ice. Cell lysates were centrifuged at 2,300 x g for 5 min at 4°C to isolate nuclei. Nuclei were suspended in 500 μl of RIPA nuclear lysis buffer [50 mM Tris-HCl (pH 7.5), 150 mM NaCl, 0.5% SDS, 0.125% sodium deoxycholate and 1% Triton X-100 in Millipore water] with 1x freshly supplemented Halt Protease Inhibitor and subjected to sonication to shear chromatin fragments to an average size of 200-500 bp on a Diagenode Bioruptor with 30s on/off cycles at high intensity for 15 minutes. Fragmented chromatin was centrifuged at 16,100 x g for 10 min at 4°C. 450 μl of supernatant was transferred to a new microcentrifuge tube. 4 μg anti-mCherry antibody (Thermo PA5-34974) was added to each sample and incubated at 4°C for 3h. 50μl of blocked Protein G Dynabeads (Thermo 10003D) was added to each sample and rotated at 4°C overnight. After overnight incubation, Dynabeads were washed seven times as described above for selection of biotinylated proteins. Chromatin was eluted from Dynabeads in 200μl elution buffer [50 mM Tris-HCl (pH 8.0), 10 mM EDTA, 1% SDS] and transferred to a new microcentrifuge tube. Eluted chromatin was treated with 1 μl RNase A and incubated overnight at 65°C to reverse crosslinks. 7.5 μl of 20 mg/ml proteinase K was added to each sample followed by incubation for 2h at 50°C. ChIP DNA was then incubated with 1ml Buffer PB (QIAGEN) and 10 μl of 3M sodium acetate pH 5.2 at 37°C for 30 minutes. DNA was purified using QIAGEN quickspin column.

15 ng of ChIP DNA was processed for library preparation using the NEBNext ChIP-seq Library Prep Kit (New England Biolabs) according to the manufacturer’s protocol.

15 ng of ChIP DNA was end-repaired using NEBNext End Repair module (NEB Cat. E6050) and purified with 1.8x AMPure XP beads (Beckman-Coulter Cat. A63880). End-repaired DNA was processed in a dA-tailing reaction using NEBNext dA-Tailing module (NEB Cat. E6053) and purified with 1.8x AMPure XP beads. Adaptor oligos 1 (5′-pGAT CGG AAG AGC ACA CGT CT-3′) and 2 (5′-ACA CTC TTT CCC TAC ACG ACG CTC TTC CGA TCT-3) used in Y-shaped adapter mix were ligated to dA-tailed DNA according to ref. ^33^ and purified with 1.5x AMPure XP beads. Ligated DNA was incubated in a thermal cycler (98°C for 40s, 65°C for 30s, and 72°C 30s) with Illumina barcode primers 2-1 (5′-CAA GCA GAA GAC GGC ATA CGA GAT CGT GATGTG ACT GGA GTT CAG ACG TGT GCT CTT CCG ATC T-3′), 2-2 (5′-CAA GCA GAA GAC GGC ATA CGA GAT ACA TCGGTG ACT GGA GTT CAG ACG TGT GCT CTT CCG ATC T-3′), 2-3′(5′-CAA GCA GAA GAC GGC ATA CGA GAT GCC TAAGTG ACT GGA GTT CAG ACG TGT GCT CTT CCG ATC T-3′) and NEB Q5 Polymerase Master Mix. Primer 1 (5′-AAT GAT ACG GCG ACC ACC GAG ATC TAC ACT CTT TCC CTA CAC GAC GCT CTT CCG ATC T-3′) was added to mix for 10 cycles (98°C for 10s, 65°C for 30s, 72°C for 30s), followed by incubation at 72°C for 3′minutes. PCR-enriched DNA was purified with 1x AMPure XP beads.

Raw Illumina sequencing reads of 150 nucleotide length were processed as fastq files in R. Reads were trimmed using the Bioconductor ShortRead R package at positions which contained 2 nucleotides in a 5-nucleotide bin with a quality encoding less than phred score = 20. Reads with at least one (TTAGGG)4 or (CCCTAA)4 segment constituted a “hit”, and were counted using the Bioconductor Biostrings R package. (number of hits / total trimmed reads) was calculated to assess the specificity of Cas9-mCherry-APEX2 for each sample.

### SILAC labeling

On day 0, early-passage, sorted, stably transduced sgTelo or sgAlpha U2OS cells were grown in heavy SILAC media, which contained L-arginine-^13^C_6_, ^15^N_4_ (Arg10) and L-lysine-^13^C_6_, ^15^N_2_ (Lys8) (Sigma). Stable sgNS cells were grown in medium SILAC media, which contained L-arginine-^13^C_6_ (Arg6) and L-lysine-4,4,5,5-*d*_4_ (Lys4) (Sigma). Untransduced U2OS cells were grown in light SILAC media, which contained L-arginine (Arg0) and L-lysine (Lys0) (Sigma). Cells were grown for more than 10 days (>5 passages) to allow for sufficient incorporation of the isotopes. On day 11, dox and Shield1 were added to each isotope culture (4 plates for each cell line) 21h before BP and H_2_O_2_ treatment. The biotinylation, nuclei isolation, and cell lysis followed the procedure described above. Before streptavidin affinity purification, equal amount of proteins measured by Pierce™ BCA Protein Assay Kit (~ 1mg from each isotope sample) were mixed in a 1:1:1 ratio (H:M:L). Streptavidin affinity purification and sample wash were described above. Proteins were eluted in 50 μl 3x protein loading buffer supplemented with 2 mM biotin and 20 mM DTT with heating for 10min at 65°C. 50 μl eluents were loaded and run approximately to the center of the lane on a 4-12% SDS-PAGE gel (Bio-Rad). The coomassie-stained protein bands were excised and cut to five slices for in-gel digestion and LC-MS/MS analysis.

### LC-MS/MS and proteomic analyses for LFQ

Unresolved protein bands from SDS-PAGE were cut into 1x1 mm pieces and placed in 1.5ml Eppendorf tubes with 1ml of water. After 30 min, water was removed and replaced with 70 μl of 250 mM ammonium bicarbonate. Proteins were then reduced by the addition of 20 μl of 45 mM 1,4-dithiothreitol, incubated at 50°C for 30 min, cooled to room temperature, alkylated with 20 μl of 100 mM iodoacetamide for 30 min, and washed twice with 1 ml water. The water was removed and replaced with 1 ml of 50 mM ammonium bicarbonate: acetonitrile (1:1) and incubated at room temperature for 1 hr. The solvent was then replaced with 200 μl acetonitrile, removed, and the pieces dried in a Speed Vac. Gel pieces were then rehydrated in 75 μl of 4 ng/μl sequencing-grade trypsin (Promega) in 0.01% ProteaseMAX Surfactant (Promega) in 50 mM ammonium bicarbonate and incubated at 37°C for 21 hr. The supernatant was then removed to a 1.5 ml Eppendorf tube, the gel pieces further dehydrated with 100 μl of acetonitrile: 1% (v/v) formic acid (4:1), and the combined supernatants dried on a Speed Vac. Peptides were then reconstituted in 25 μl of 5% acetonitrile containing 0.1% (v/v) trifluoroacetic acid for LC-MS/MS.

Samples were analyzed on a NanoAcquity UPLC (Waters Corporation) coupled to a Q Exactive (Thermo Fisher Scientific) hybrid mass spectrometer. In brief, 1.0 μl aliquots were loaded at 4 μl/min onto a custom-packed fused silica precolumn (100 μm ID) with Kasil frit containing 2 cm Magic C18AQ (5μm, 100A) particles (Bruker Corporation). Peptides were then separated on a 75μm ID fused silica analytical column containing 25 cm Magic C18AQ (3μ, 100A) particles (Bruker) packed in-house into a gravity-pulled tip. Peptides were eluted at 300 nl/min with a linear gradient from 95% solvent A (0.1% (v/v) formic acid in water) to 35% solvent B (0.1% (v/v) formic acid in acetonitrile) in 60 min. Data was acquired by data-dependent acquisition according to a published method^34^. Briefly, MS scans were acquired from *m/z* 300-1750 at a resolution of 70,000 (*m/z* 200) and followed by ten tandem mass spectrometry scans using HCD fragmentation using an isolation width of 1.6 Da, a collision energy of 27%, and a resolution of 17,500 (*m/z* 200). Raw data files were processed with Proteome Discoverer (Thermo, version 2.1.1.21) and searched with Mascot (Matrix Science, version 2.6) against the SwissProt *Homo sapiens* database. Search parameters used tryptic specificity considering up to 2 missed cleavages, a parent mass tolerance of 10 ppm, and a fragment mass tolerance of 0.05 Da. Fixed modification of carbamidomethyl cysteine was considered as were variable modifications of N-terminal acetylation, N-terminal conversion of Gln to pyroGlu, oxidation of methionine, and biotin-phenol conjugation of tyrosine. Results were loaded into Scaffold (Proteome Software Inc., version 4.8.4) for peptide and protein validation and quantitation using the Peptide Prophet and Protein Prophet algorithms^35^’ ^36^. The threshold for peptides was set to 80% (1.1% FDR) and 90% for proteins (3-peptide minimum). Contaminants such as human keratin were included in all statistical analyses and removed from the figures.

### LC-MS/MS and proteomic analyses for SILAC

A fully resolved SDS-PAGE was cut into 5 fractions and each fraction was processed separately as described. Gel bands were cut into 1x1 mm pieces and placed in 1.5 mL eppendorf tubes with 1mL of water for 30 min. The water was removed and 200μl of 250 mM ammonium bicarbonate was added. For reduction, 20 μl of a 45 mM solution of 1,4-dithiothreitol (DTT) was added and the samples were incubated at 50°C for 30 min. The samples were cooled to room temperature and then, for alkylation, 20 μl of a 100 mM iodoacetamide solution was added and allowed to react for 30 min. The gel slices were washed twice with 1 mL water. The water was removed and 1mL of 50:50 (50 mM ammonium bicarbonate:acetonitrile) was placed in each tube and samples were incubated at room temperature for 1h. The solution was then removed and 200 μl of acetonitrile was added to each tube, at which point the gels slices turned opaque white. The acetonitrile was removed and gel slices were further dried in a Speed Vac (Savant Instruments, Inc.). Gel slices were rehydrated in 100 μl of 4ng/μl of sequencing-grade trypsin (Sigma) in 0.01% ProteaseMAX Surfactant (Promega):50 mM ammonium bicarbonate. Additional bicarbonate buffer was added to ensure complete submersion of the gel slices. Samples were incubated at 37°C for 18 hrs. The supernatant of each sample was then removed and placed in a separate 1.5 mL eppendorf tube. Gel slices were further extracted with 200 μl of 80:20 (acetonitrile:1% formic acid). The extracts were combined with the supernatants of each sample. The samples were then completely dried down in a Speed Vac.

Tryptic peptide digests were reconstituted in 25 μl 5% acetonitrile containing 0.1% (v/v) trifluoroacetic acid and separated on a NanoAcquity (Waters) UPLC. In brief, a 3.0 μl injection was loaded in 5% acetonitrile containing 0.1% formic acid at 4.0 μL/min for 4.0 min onto a 100 μm I.D. fused-silica pre-column packed with 2 cm of 5 μm (200Å) Magic C18AQ (Bruker-Michrom) and eluted using a gradient at 300 nL/min onto a 75 μm I.D. analytical column packed with 25 cm of 3′μm (100Å) Magic C18AQ particles to a gravity-pulled tip. The solvents were A, water (0.1% formic acid); and B, acetonitrile (0.1% formic acid). A linear gradient was developed from 5% solvent A to 35% solvent B in 60 minutes. Ions were introduced by positive electrospray ionization via liquid junction into a Q Exactive hybrid mass spectrometer (Thermo). Mass spectra were acquired over *m/z* 300-1750 at 70,000 resolution (*m/z* 200) and data-dependent acquisition selected the top 10 most abundant precursor ions for tandem mass spectrometry by HCD fragmentation using an isolation width of 1.6 Da, collision energy of 27, and a resolution of 17,500.

Raw data files were peak processed with Mascot Distiller (Matrix Science, version 2.6) prior to database searching with Mascot Server (version 2.6) against the *Uniprot_Human* database. Search parameters included trypsin specificity with two missed cleavages. The variable modifications of oxidized methionine, pyroglutamic acid for N-terminal glutamine, N-terminal acetylation of the protein, biotin-phenol on tyrosine and a fixed modification for carbamidomethyl cysteine were considered. For SILAC labels, the medium samples were labeled with Lys4 and Arg6 and the heavy samples were labeled with Lys8 and Arg10. The mass tolerances were 10 ppm for the precursor and 0.05 Da for the fragments. SILAC ratio quantitation was accomplished using Mascot Distiller and the results from Mascot Distiller were loaded into the Scaffold Viewer (Proteome Software, Inc., version 4.8.4) for peptide/protein validation and SILAC label quantitation. For SILAC experiments, protein identification was subject to a two-peptide cut-off. For proteins detectable in the H sample but that lack an empirical H/L ratio value (due to low background detection in the L sample), peak areas of all the identified peptides in the Distiller file were used to calculate H/L ratios.

### Data analysis

Data was first filtered to exclude proteins detected in only one of the dCas9-mCherry-APEX2/sgTelo (+BP, +H_2_O_2_) (“S1”) replicates, followed by log_2_ transformation. Prior to the log_2_ transformation, iBAQ values of 0 were replaced with the smallest iBAQ value from the corresponding sample in dCas9-mCherry-APEX2/sgTelo (+BP, ‐H_2_O_2_) (“S2”) or dCas9-mCherry-APEX2/sgNS (+BP, +H_2_O_2_) (“S3”) to avoid generation of infinite ratios. Moderated t-test with a paired design was used to compare the log_2_ transformed iBAQ values between S1 and S3, S1 and S2, and S2 and S3 using limma package^37^. To adjust for multiple comparisons, *p* values were adjusted using the Benjamini-Hochberg (BH) method^38^. Proteins were selected for subsequent analysis if they were (*i*) significantly enriched in both S1 vs. S3 and S1 vs. S2, (*ii*) not enriched in S2 vs. S3, and (*iii*) if S1/S3 and S1/S2 ratios were greater than 2.

Similarly, SILAC datasets were filtered to exclude proteins with H/M ratios detected in only one of the biological replicates. Detection in a biological replicate required identification in at least two of the three technical replicates that were done for each biological replicate; median values from the technical replicates were used for subsequent analyses. Proteins with BH-adjusted *p* values less than 0.01 (moderated t-test described above) are considered statistically significant. Proteins (with BH-adjusted *p* values < 0.01 and log_2_ fold change ≥ 2.5) were selected for subsequent GO (David Bioinformatics) and overlap analysis. To determine whether the proteins identified in this experiment overlap significantly with three published datasets, hypergeometric test was used. Hypergeometric test was also used for testing the overlapping proteins between C-BERST telomere IDs and centromere IDs.

---

**Supplementary Fig. 1.**
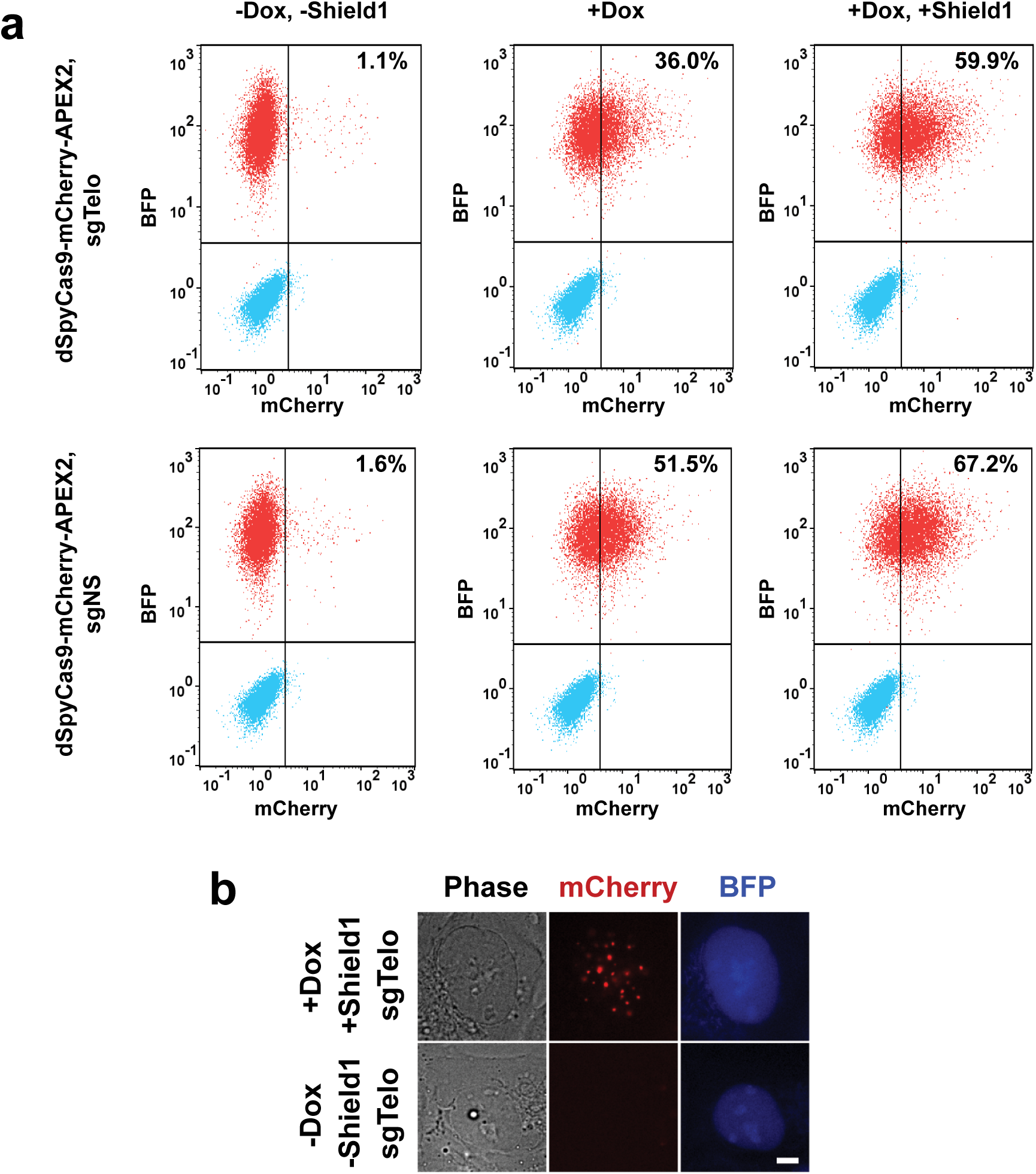
Inducible dSpyCas9-mCherry-APEX2 expression. **(a)** Flow cytometry was used to measure the percentage of mCherry+ and BFP+ double-positive cells under different induction conditions. Stable U2OS cells expressing sgTelo (top row) or sgNS (bottom row) were exposed to three conditions for 21h before flow cytometry: no inducers (left), dox only (2 μg/ml, middle), or a combination of dox (2 μg/ml) and Shield1 (250 nM) (right). Cyan: untransduced cells; red: transduced cells. With both sgRNAs, dox and Shield1 in combination yield the highest percentages of double-positive cells. Specific percentages of mCherry+, BFP+ cells are indicated in each plot. **(b)** Live-cell imaging of clonal cells derived from the sgTelo P1 population (see **Fig. 1c**). When inducers are omitted, dSpyCas9-mCherry-APEX2 expression and telomeric accumulation are not observed. Scale bar, 5 μm.

**Supplementary Fig. 2.**
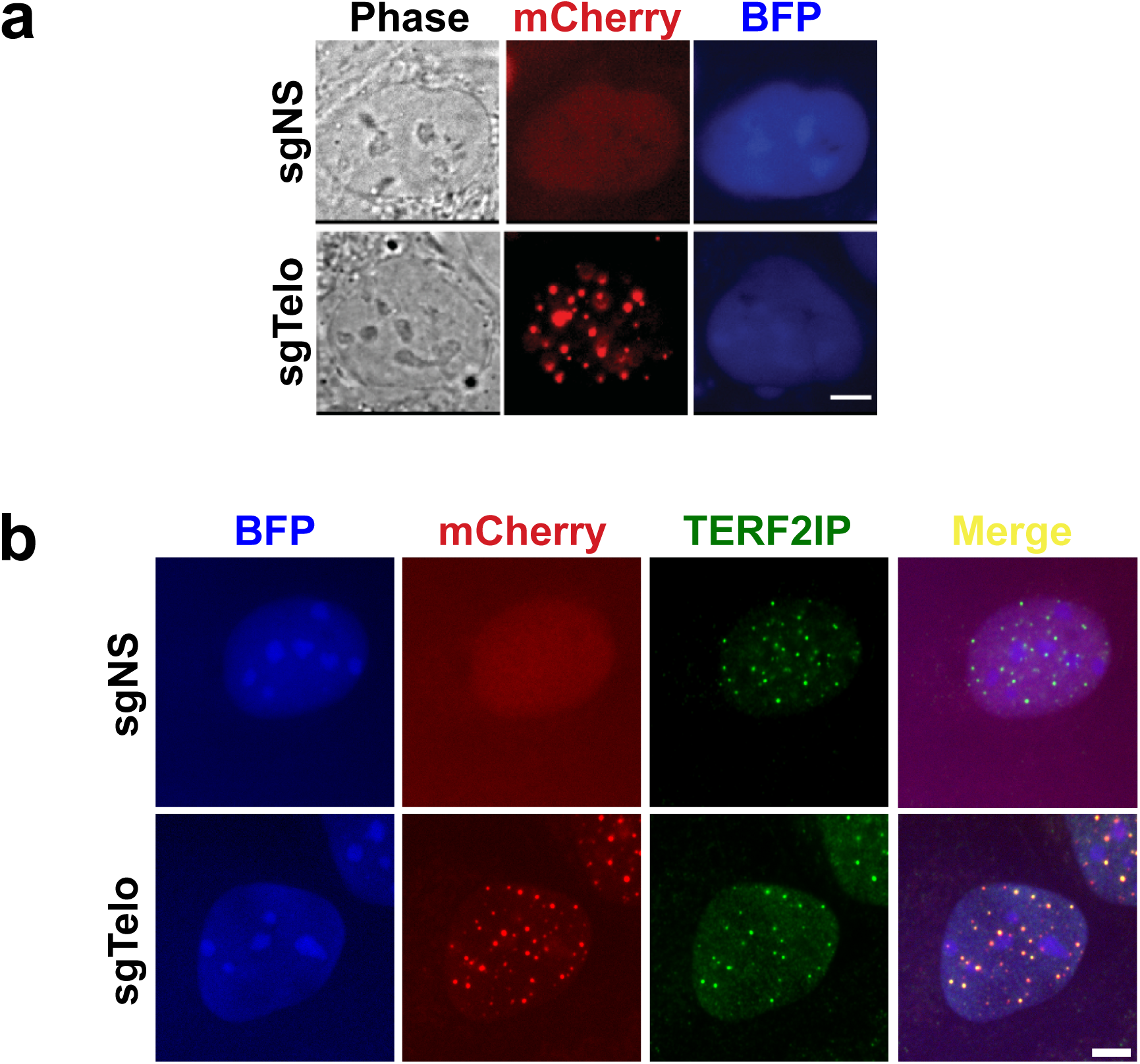
Specific telomere targeting by dSpyCas9-mCherry-APEX2. **(a)** Livecell imaging of telomere localization by dSpyCas9-mCherry-APEX2 in U2OS cells, using the P1-sorted population defined by the FACS workflow in **Fig. 1c**. dSpyCas9-mCherry-APEX2 exhibited telomeric foci with sgTelo but not with sgNS. **(b)** Immunostaining of telomeric marker protein with primary anti-TERF2IP and secondary antibody conjugated with Alexa 488. Colocalization of dSpyCas9-mCherry-APEX2 foci with TERF2IP is observed (n ≥ 25 cells examined). Scale bar, 5 μm.

**Supplementary Fig. 3.**
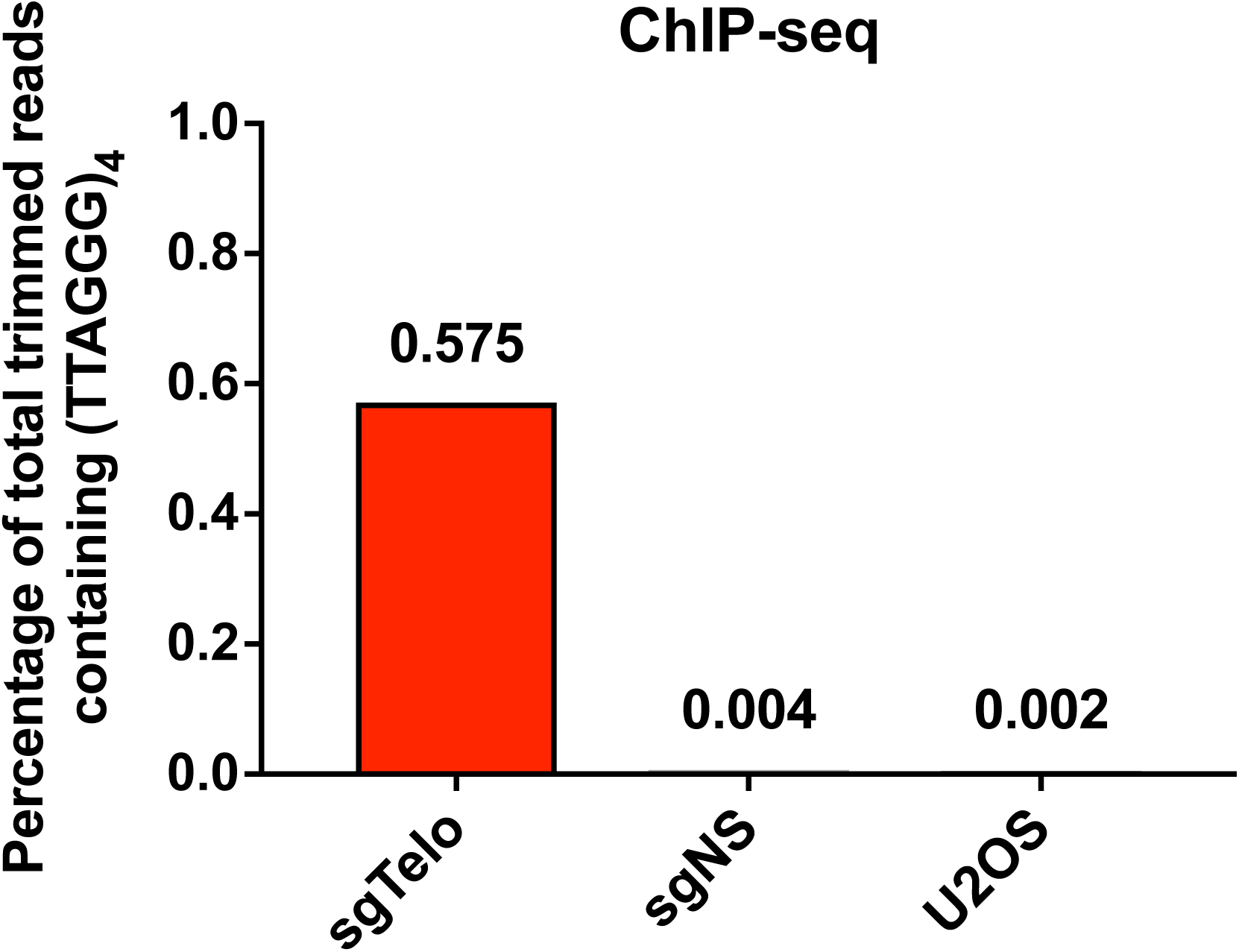
anti-mCherry chromatin immunoprecipitation shows genome-wide binding of sgTelo-programmed dSpyCas9-mCherry-APEX2. The reads were trimmed by adaptor removal and filtering. The percentage of total trimmed reads that include at least one (TTAGGG)_4_ telomeric sequence (the minimum length required for complete sgTelo complementarity) is shown. Values were averaged from two independent biological replicates, except for the U2OS ChIP-seq, which was only performed once.

**Supplementary Fig. 4.**
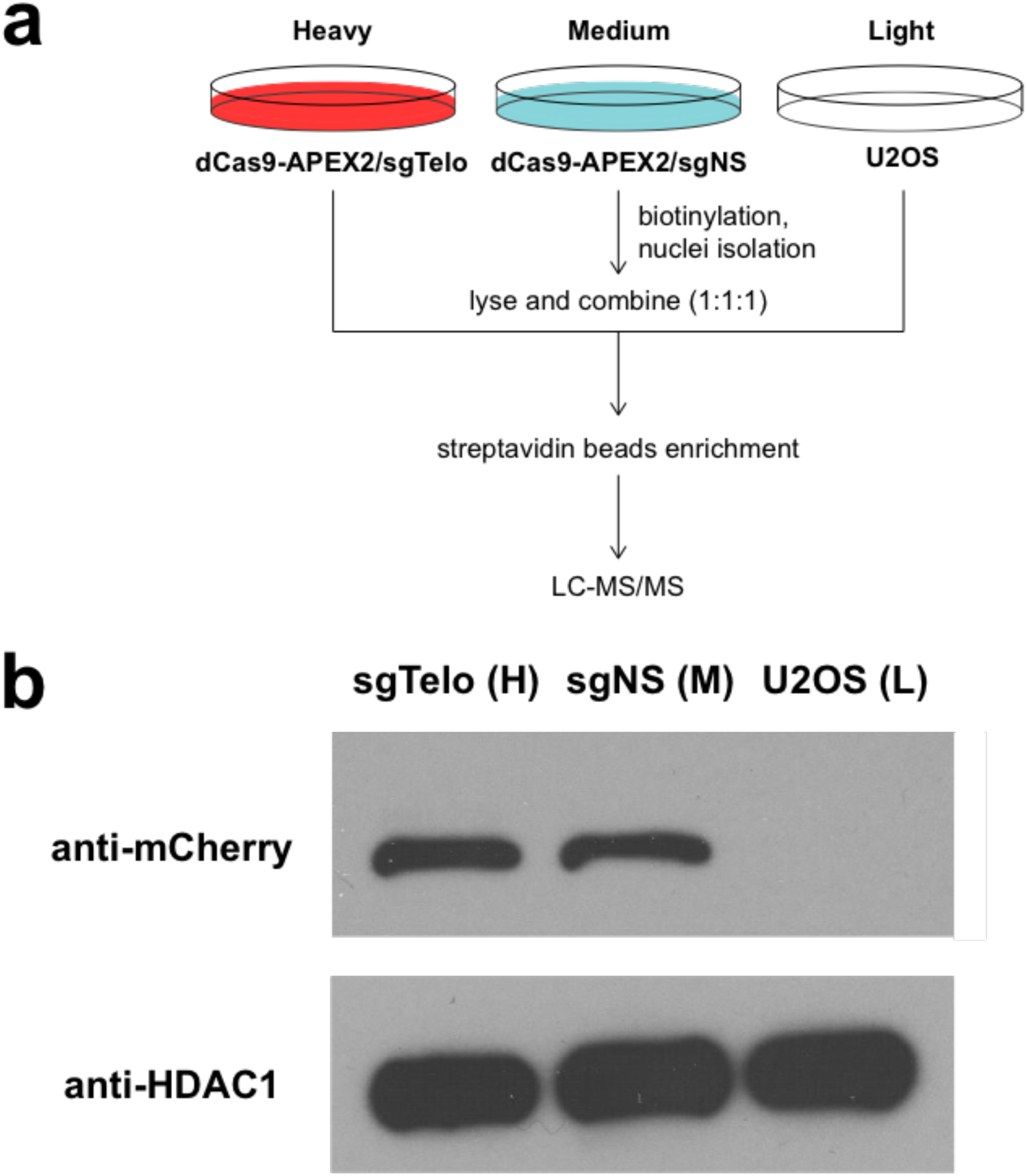
**(a)** Schematic diagram of SILAC workflow. Cells were grown in different isotope culture media for at least five passages. dSpyCas9-mCherry-APEX2 proteins were induced by dox and Shield1 21 hours before biotinylation. Following biotinylation and nuclei isolation, cell lysates were sonicated and mixed in a 1:1:1 ratio. **(b)** Anti-mCherry was used to detect dSpyCas9-mCherry-APEX2 (top), and anti-HDAC1 was used as a loading control (bottom) in the western blot analysis.

**Supplementary Fig. 5.**
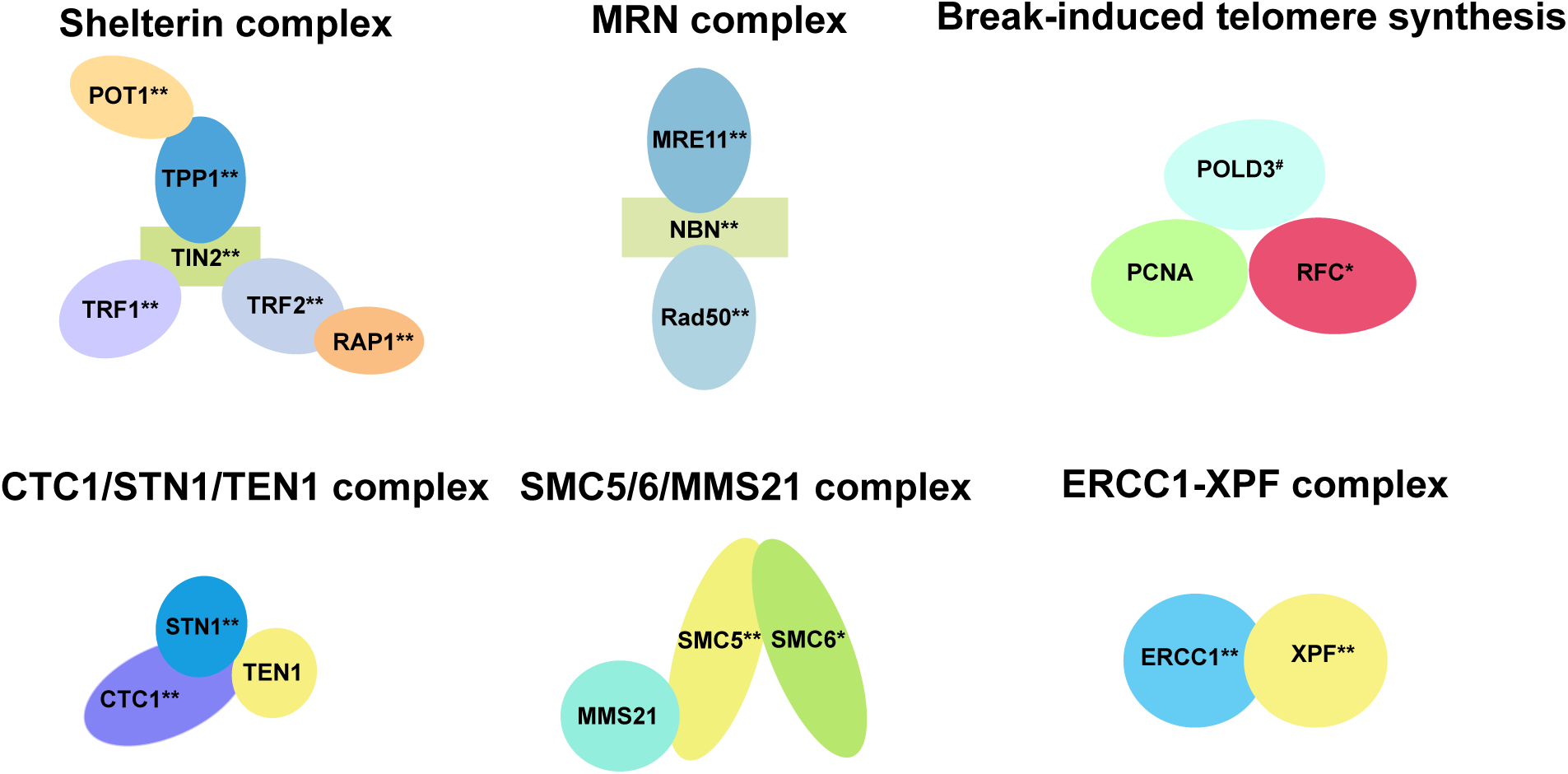
C-BERST specifically detects components of multiple complexes and factors implicated in ALT pathways. Proteins denoted by double asterisks were significantly enriched ( *p* value < 0.01) and meet the SILAC cut-off log_2_ fold change ≥ 2.5 in the sgTelo labeling sample, relative to the sgNS labeling samples (H/M) ratio (see **Supplementary Table 3**). Proteins denoted by single asterisks were also detected but with lower degrees of significance (BH-adjusted *p* value < 0.05) in the sgTelo labeling sample (**Supplementary Table 3**). Proteins denoted by a hashtag were enriched and statistically significant in LFQ. Components of the RAD9/RAD1/HUS1 complex (see **Supplementary Table 3**) were not detected.

**Supplementary Fig. 6.**
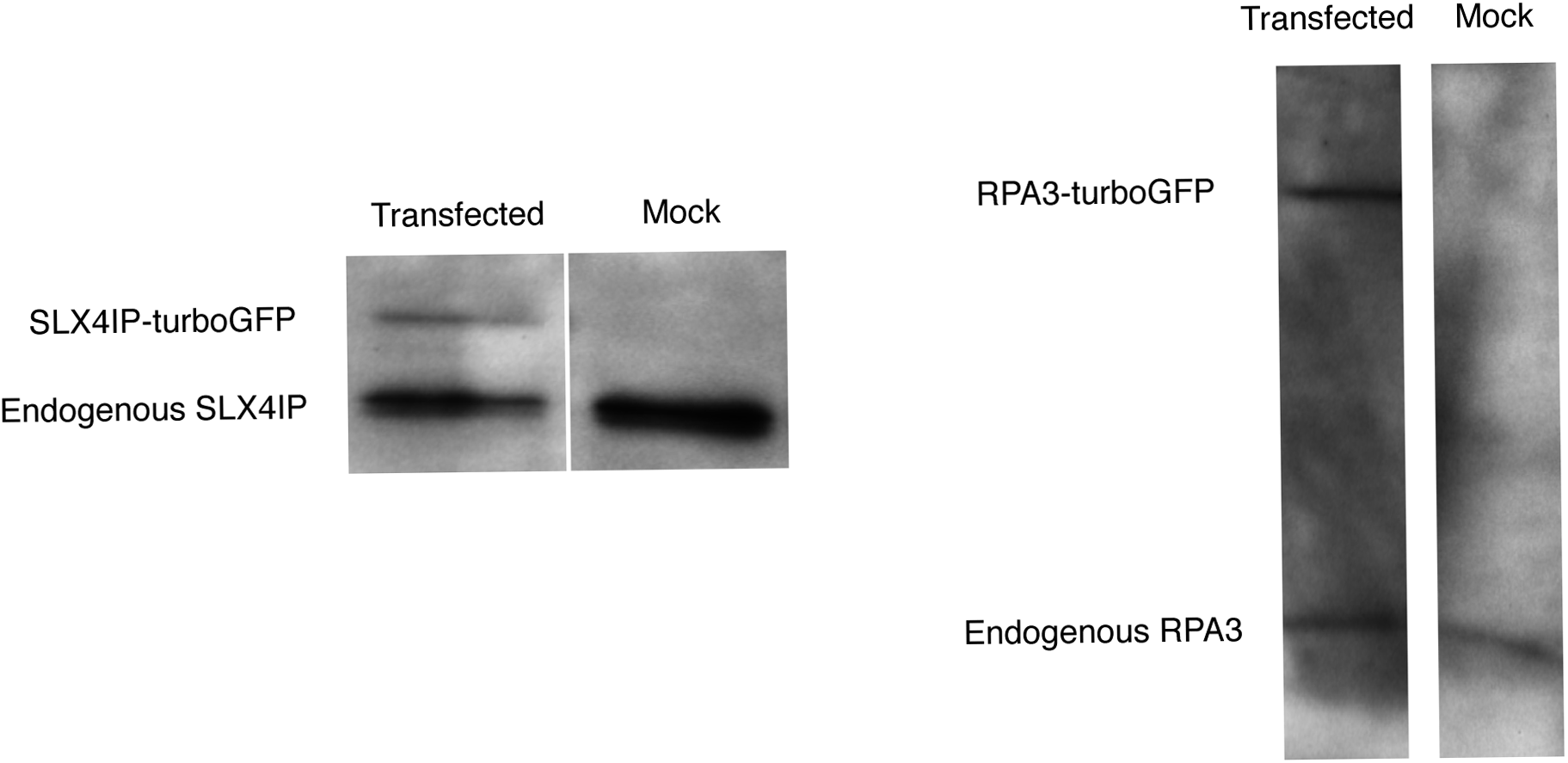
Western blot analysis of exogenous, turboGFP-tagged SLX4IP or RPA3 expression, in comparison with the corresponding endogenous protein. ~0.1 x 10^5^ U2OS cells transfected with SLX4IP-turboGFP or RPA3-turboGFP expression plasmid (100 ng and 50 ng, respectively) were lysed in 1x RIPA lysis buffer, and proteins were resolved by SDS-PAGE. Western blots were probed by primary SLX4IP or RPA3 antibody and anti-rabbit secondary antibody conjugated with HRP. The gel lanes shown in each panel were cropped from identical exposures of the same western blot membranes.

**Supplementary Fig. 7.**
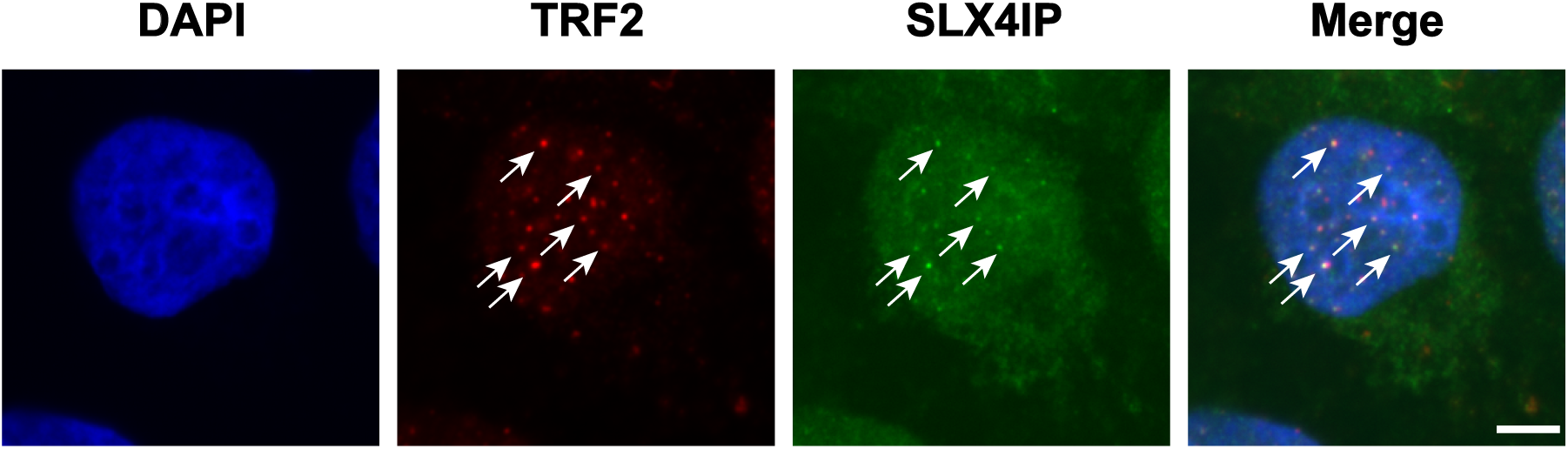
Coimmunostaining of TRF2 (telomeric marker protein) and SLX4IP protein. Primary goat anti-TRF2 and rabbit anti-SLX4IP were used to detect endogenous TRF2 and SLX4IP in the fixed U2OS cells. Secondary donkey anti-goat conjugated with Alexa 647 and mouse anti-rabbit conjugated with CruzFluor^TM^ 488 were then incubated with cells. Scale bar, 5 μm.

**Supplementary Fig. 8.**
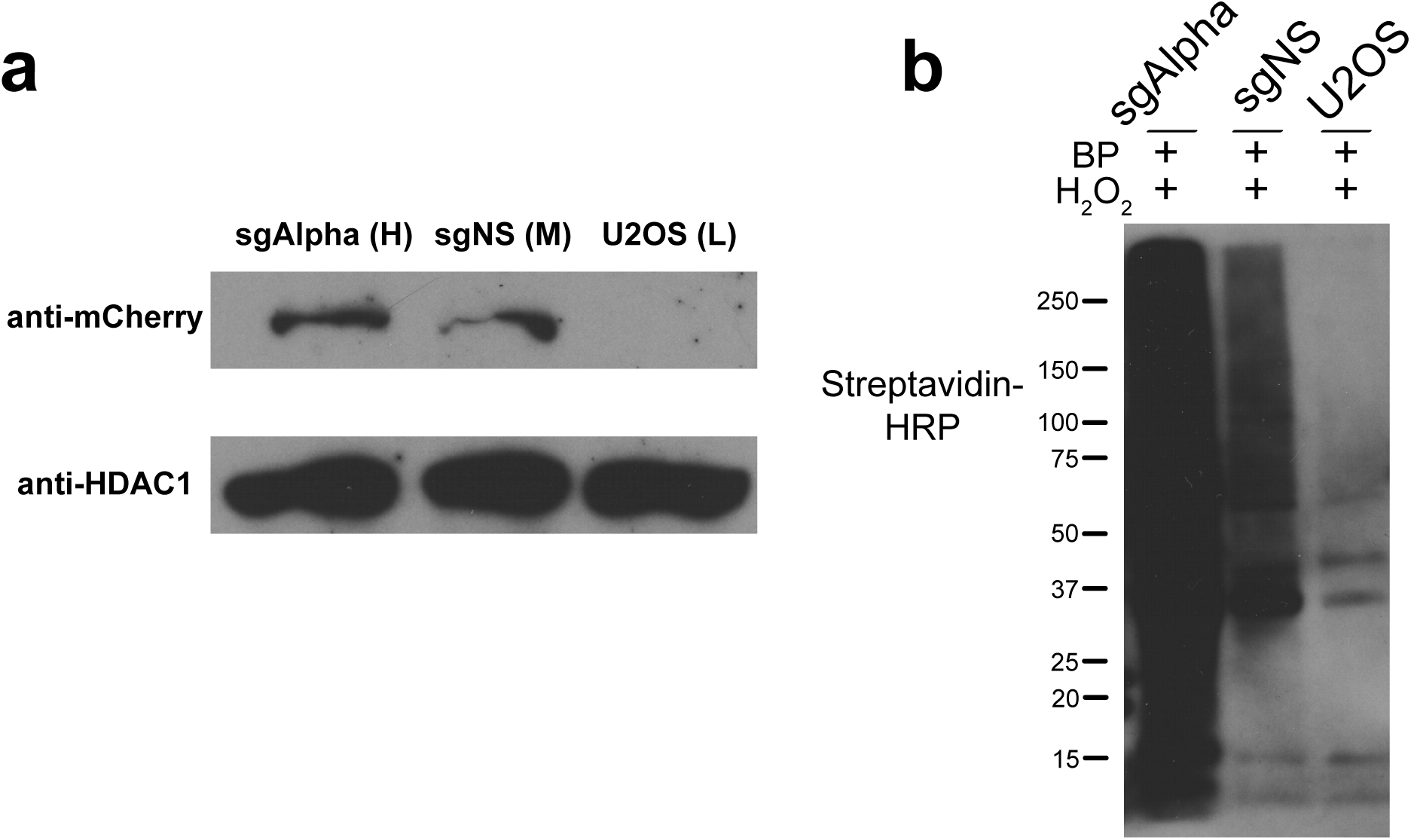
**(a)** Western blot analysis of dSpyCas9-mCherry-APEX2 expression using anti-mCherry to detect dSpyCas9-mCherry-APEX2 (top). Anti-HDAC1 was used as a loading control (bottom). **(b)** Streptavidin-HRP blotting analysis of biotinylated proteins in sgAlpha‐ and sgNS-expressing cells. Untransduced U2OS cells were used as a control.

**Supplementary Fig. 9.**
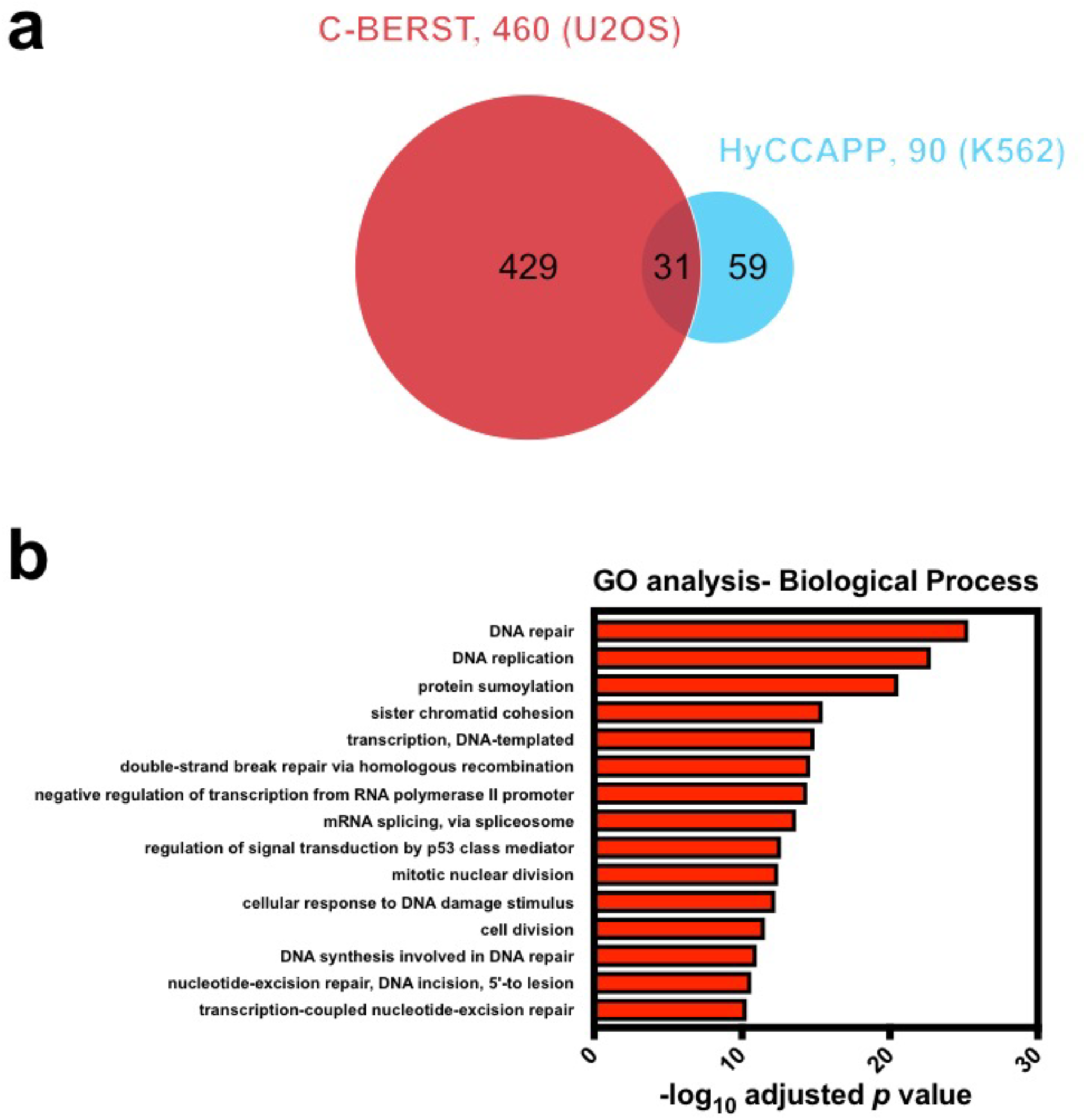
**(a)** Venn diagram of statistically enriched (BH-adjusted *p* value < 0.01) 460 centromeric proteins from U2OS cells, as detected by C-BERST (red), and 90 centromeric proteins from K562 cells, as detected by HyCCAPP (cyan) (see text). 31 proteins from the C-BERST proteome were detected by both. **(b)** Gene Ontology-Biological Process (GO-BP) analysis on 460 centromeric proteins identified by C-BERST. The x-axis is the ‐log_10_ *p* value (BH-adjusted) for the C-BERST-detected proteins associated with each GO term given on the left. The 15 most statistically significant GO terms are displayed.

**Supplementary Fig. 10.**
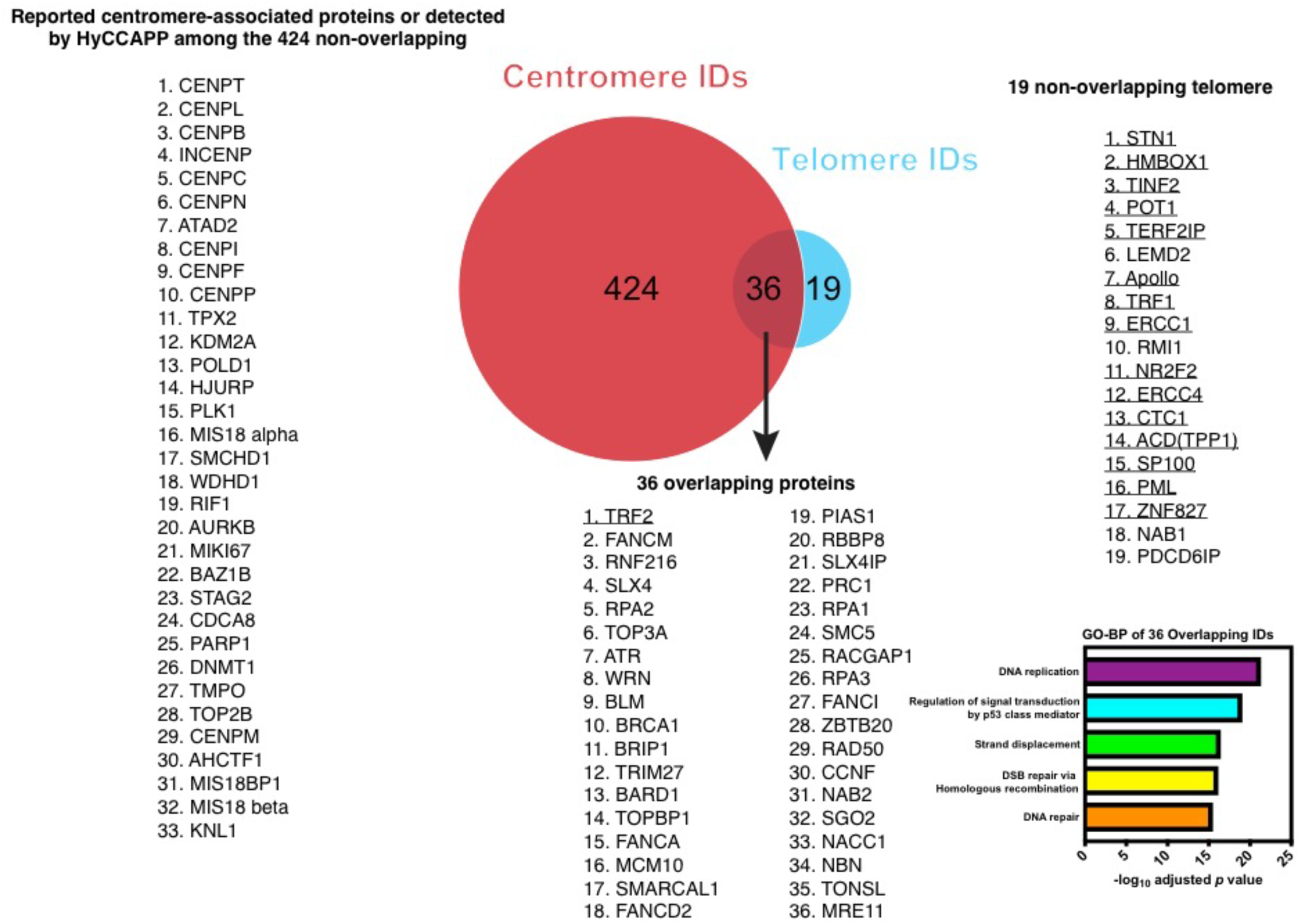
Venn diagram of 55 C-BERST ALT/telomeric IDs and 460 centromeric IDs. 19 non-overlapping telomere IDs are listed on the right. Known ALT/telomeric proteins are underlined. Among the 424 non-overlapping centromere IDs, 33 (listed on the left) are known or implicated as centromeric proteins. 36 overlapping proteins from both sets are listed below the Venn diagram, as indicated. The five most significant GO-BP terms for the 36 overlapping ID are provided on the lower right.

## Supplementary Note

SpyCas9-mCherry-APEX2, sgTelo, and sgNS sequences.

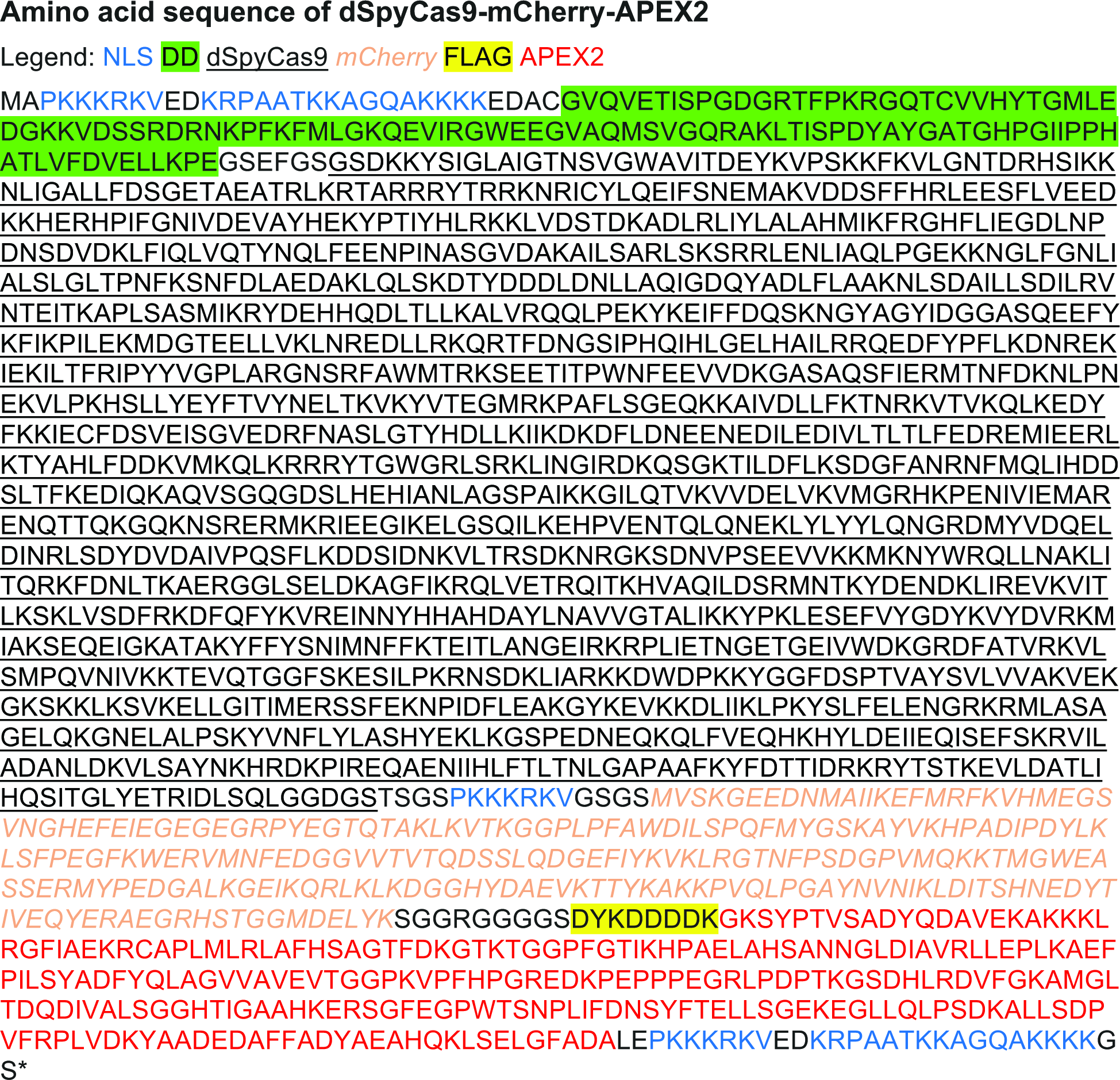

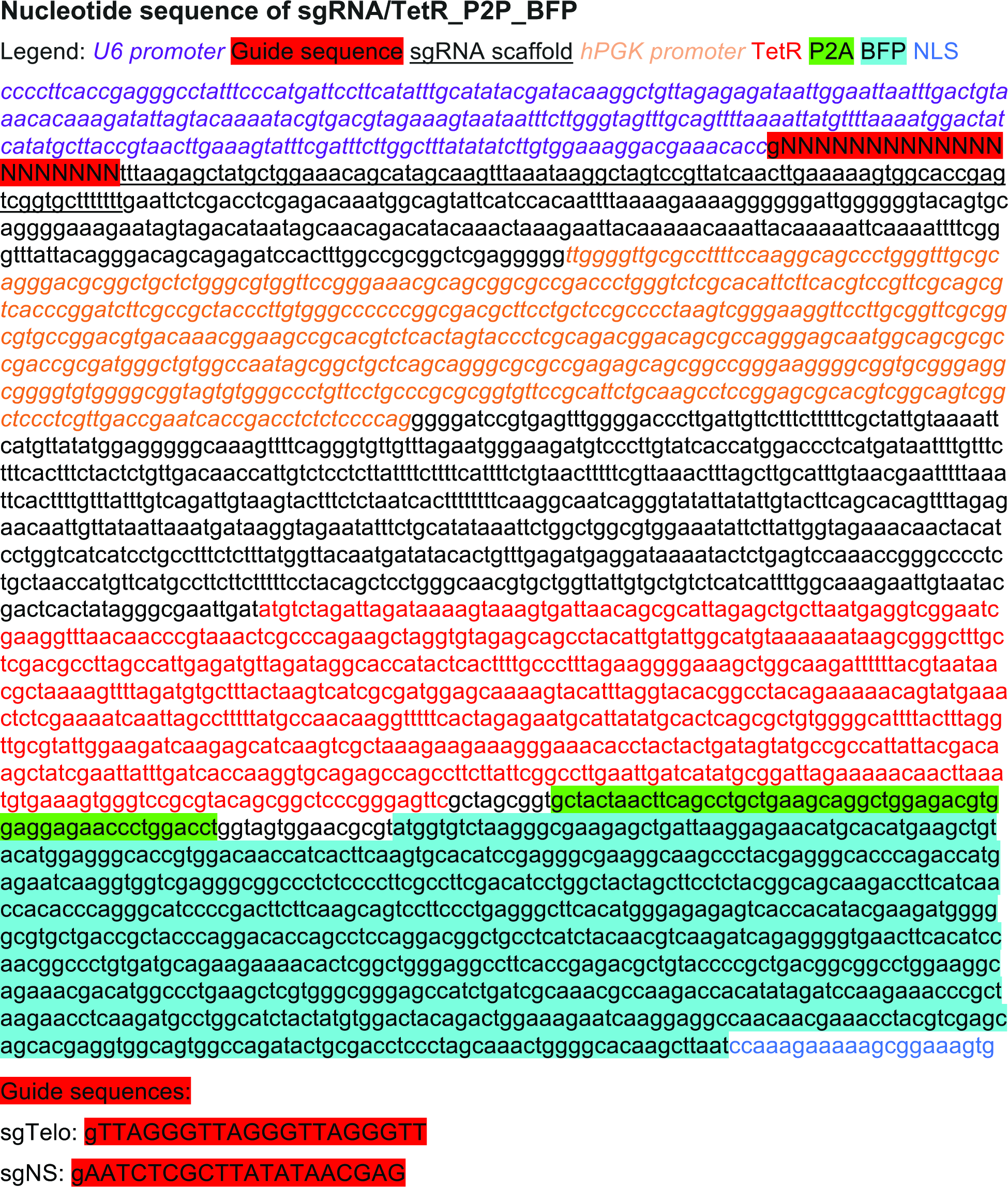

